# Simple and thorough detection of related sequences with position-varying probabilities of substitutions, insertions, and deletions

**DOI:** 10.1101/2025.03.14.643233

**Authors:** Martin C. Frith

## Abstract

One way to understand biology is by finding genetic sequences that are related to each other. Often, a family of related sequences has position-varying probabilities of substitutions, insertions, and deletions: we can use these to find distantly-related sequences. There are popular software tools for this task, which all have limitations. They either do not use all probability evidence (e.g. PSI-BLAST, MMseqs2), or have excessive complexity and minor biases (e.g. HMMER). This complexity inhibits fertile development of alternative tools.

This study describes a simplest reasonable way to find related sequences, making full use of position-varying probabilities. The algorithms likely use the fewest operations that such algorithms possibly could, so they are fast and simple. This has been implemented in prototype software named DUMMER (Dumb Uncomplicated Match ModelER). Its sensitivity and specificity are competitive with HMMER. It finds evidence that the human genome has vastly more relics of LF-SINE, retrotransposons that were co-opted for various functions in common ancestors of all land vertebrates.

## Introduction

This study was motivated by seeking subtly-related DNA sequences. Non-coding DNA is hardly ever conserved between vertebrates and invertebrates, but we recently found gene-regulating DNA conserved between humans, molluscs, arthropods, and even corals (Frith and Ni 2023). These DNA regions all control gene expression in embryonic development: a control system conserved since early Precambrian animals.

These DNA sequences have position-varying rates of substitutions, insertions, and deletions, for example, a conserved “CCAAT box” near the right of Fig. 1. This suggests we could find more-subtly related sequences by using these position-specific probabilities. This is a classic approach: a set of position-specific rates is termed a “profile”, and there are many methods for comparing a profile to a sequence, to find similar regions between the sequence and the profile (Gribskov et al. 1987, Krogh et al. 1994, Karplus et al. 1998, Yu et al. 2002, Wheeler and Eddy 2013, Steinegger and Söding 2017).

**Figure 1.**
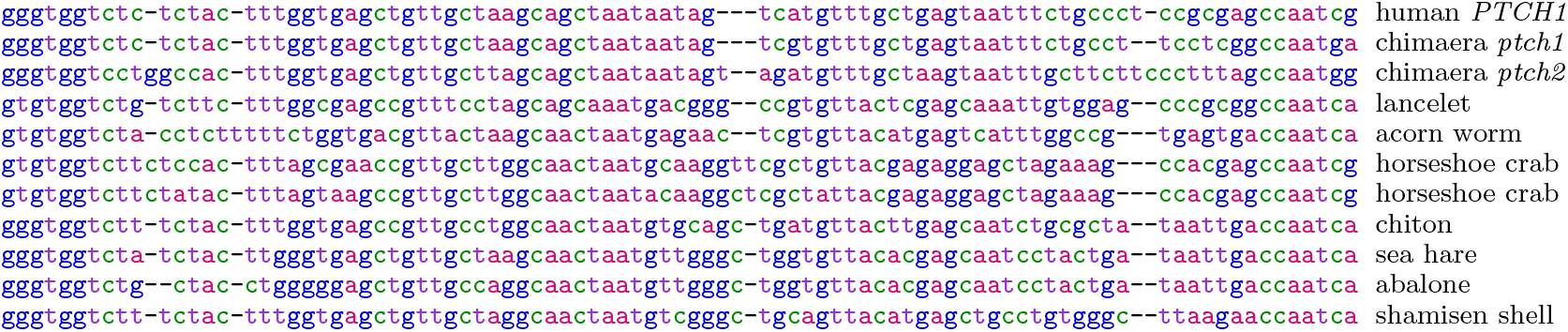
Alignment between promoter DNA sequences of *Ptch* (patched) genes in some animals (Frith and Ni 2023).

We can seek “local” similarities between part of a profile and part of a sequence, or “semi-global” similarities between a whole profile and part of a sequence. The popular tools are all local. The main reason is that it’s hard to determine the probability of a semi-global similarity occuring by chance in a random sequence.

The currently popular methods for comparing a profile to a sequence are of two types. The first is exemplified by PSI-BLAST: it uses position-specific rates of substitution, but its insertion and deletion parameters are position-nonspecific and ad hoc (Altschul et al. 1997). Also, it considers just one alignment between potentially related regions. It would be more powerful to combine the evidence from alternative ways of aligning them (Eddy 2008, Frith 2024).

The other type of method uses position-specific probabilities of substitution, insertion, and deletion, and combines alternative alignments. The de facto standard tool seems to be HMMER, which is complex and quirky (Eddy 2008). The perceived need for this complexity might explain why few of the many sequence comparison tools use this full-probability approach. The present study shows that this complexity is not necessary.

HMMER is designed for a question that is subtly wrong. It is designed to judge whether a sequence has regions related to a profile (Eddy 2008). This leads to complexity: it has parameters that change with the length of each sequence that is compared to a profile, so that equally good similarities get lower scores in longer sequences. This corresponds to an assumption that related regions are equally likely per sequence, rather than per unit sequence length (Frith 2024). The real aim, surely, is to find as many matches (e.g. functional domains) as possible, regardless of whether they are in long/short same/different sequences.

HMMER also strives for uniform probability of alignment start/end positions in a profile. This leads to a complex “implicit probabilistic model” (Eddy 2008). It also means that HMMER is slightly biased towards longer alignments with shorter unaligned flanks (see below).

HMMER doesn’t allow insertions next to deletions. Some earlier profile methods had position-specific probabilities for insertions next to deletions (Krogh et al. 1994). These probabilities tend to be low, so much data is needed to learn them accurately. The present study allows insertions next to deletions, without requiring extra probabilities for them.

All these tools produce a similarity score, indicating the strength of evidence for related sequences. It is important to know the probability of a similarity score occurring by chance, in a random sequence. Unfortunately, this is hard to determine. There is a solution for gapless local alignment, in the limit of infinitely long sequences (Karlin and Altschul 1990). Gapped alignment relies on conjecture and simulations, which limit PSI-BLAST to its simple profile search. HMMER uses conjectures that apply to its sophisticated profile search. The scope of HMMER’s conjectures is unclear: their practical application does not always give accurate results (Eddy 2008), and they seem to contradict the gapless solution described by Karlin and Altschul (1990) (see below).

The terms *p*-value and *E*-value are used in confusingly varying ways. This study defines them as follows. If we compare a profile to one or more random sequences:

- the *p*-value is the probability of a score ≥*s* occurring,
- the *E*-value is the expected number of distinct sequence regions with score ≥ *s*.

In a long random sequence, similarities to a profile occur randomly at a uniform rate. So the number of similar regions follows a Poisson distribution (in the limit where the sequence is long). Therefore *p* = 1 −*e*^−*E*^.

This study shows a minimal, bias-free way to find sequence parts related to a profile, using position-specific substitution, deletion, and insertion rates, and combining evidence from alternative alignments. It determines *E*-values using a conjecture of Yu and Hwa (2001). It extends a recent position-nonspecific method (Frith 2024): the non-obvious aspect of this extension is how precisely to define alignment probabilities. It doesn’t consider profile-versus-profile search, which is powerful but outside our scope (Steinegger et al. 2019).

## Methods

### Alignment scores

Let us start by reviewing score-based alignment. We wish to find related parts between a profile of length *m*, and a sequence of length *n* (*Q*_0_, *Q*_1_, …, *Q*_n−1_). The profile has these position-specific scores:

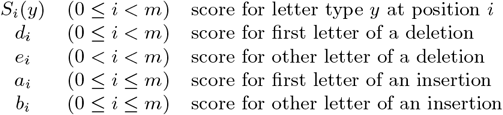

Typically, *a*_i_ < *b*_i_ < 0 and *d*_i_ < *e*_i_ < 0. This is the standard “affine gap” scheme, which works quite well because insertions and deletions are somewhat rare (low start probability) but often long (high extension probability).

Here, “insertion” means insertion in the sequence relative to the profile, and “deletion” likewise. Insertions can occur at *m* + 1 sites: between adjacent positions in the profile, or before the first position, or after the last position.

### Finding the maximum alignment score

We wish to find the maximum possible alignment score between any part of the sequence and any part of the profile. This can be found by the Smith-Waterman-Gotoh algorithm (Smith and Waterman 1981, Gotoh 1982). A slightly unusual version is shown in Algorithm 1. It considers each length-*i* prefix of the profile and length-*j* prefix of the sequence:
then removed alignments that have any

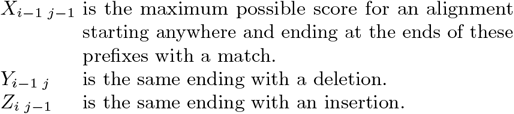

Thus, *w* is the maximum score for any alignment ending at the ends of these prefixes. Finally, the maximum score is the maximum *w*.

It might be more intuitive to write the algorithm with shifted definitions:

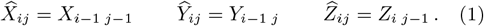

This is merely a change of notation, and it obscures the similarity to the next algorithm we will consider.

This version of the Smith-Waterman-Gotoh recurrences probably has the lowest possible number of operations: max, add, and access to previous *X, Y* and *Z* values (Cameron et al. 2004). It also has simple initialization, just setting values to −*∞*. As a tradeoff, it does excess calculations at the borders: when *i* = 0 or *m*, or *j* = 0 or *n*. In particular, it uses undefined values *e*_0_, *S*_m_, *d*_m_, *e*_m_, and *Q*_n_. They can be set to arbitrary values, and have no effect on the outcome.

So far, we just have the maximum score. We can find an alignment with this score by tracing back from a point (*i, j*) with maximum *w* (see the Supplement). This does not require storing any trace pointers in Algorithm 1.

### Limited alignment extension from an anchor

The Smith-Waterman-Gotoh algorithm is rarely used in practice, because it is too slow for large data. Instead, fast heuristics are used to find potentially related regions, which are then aligned. One way is to fix an “anchor” (tiny alignment), and extend alignments in a limited range on both sides (Fig. 2). The ranges (grey tiles) are not fixed in advance: they are adjusted as the algorithm proceeds. Several ways to do that have been suggested (Zhang et al. 1998, Suzuki and Kasahara 2017, Liu and Steinegger 2023).

**Figure 2.**
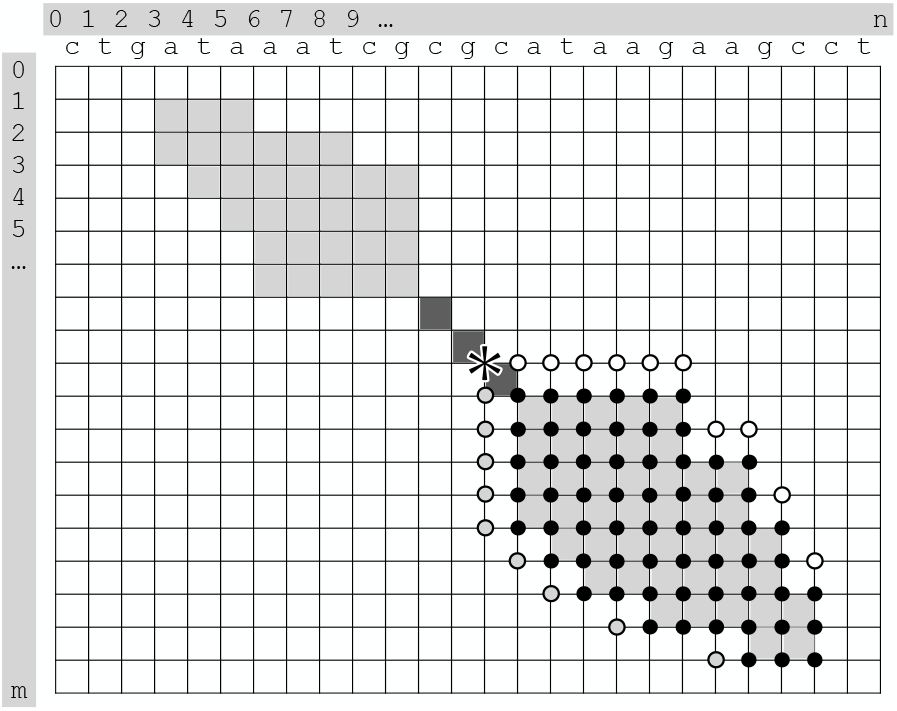
Extending alignments in a limited range either side of an anchor. The dark grey tiles show the anchor; the light grey tiles show the regions in which we seek alignments extending from it. The sequence is shown horizontally, with coordinates 0 ≤ *j* ≤ *n*. The profile is vertical, with coordinates 0 ≤ *i* ≤ *m*. The circles and star mark calculations in Algorithms 2 and 4.

This alignment extension algorithm is shown in Algorithm 2. It is almost the same as Algorithm 1: the main change is that the “0” is moved from the recurrence to the initialization at the extension start point. This is because we only consider alignments starting at this point, rather than starting anywhere.

Again, it is possible to use shifted definitions (Equations 1), but I suspect that makes it tricky to handle the irregular grey regions.

#### Algorithm 1

Maximum alignment score

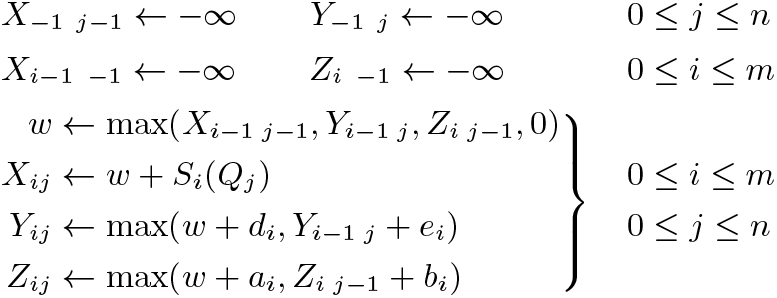

#### Algorithm 2

Maximum alignment extension score

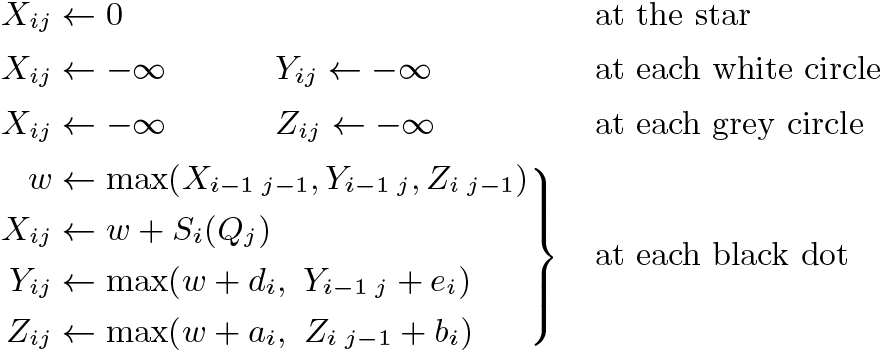

### Summing alignment probabilities

Score-based alignment is equivalent to probability-based alignment (Frith 2020). An alignment’s score is:

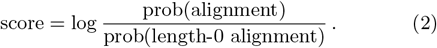

Here, prob(alignment) is the product of probabilities of insertions, deletions, aligned letters, and unaligned letters. The base of the log doesn’t matter (this study uses log_2_): changing the base just scales the scores by a constant amount.

#### Algorithm 3

Sum of alignment probabilities

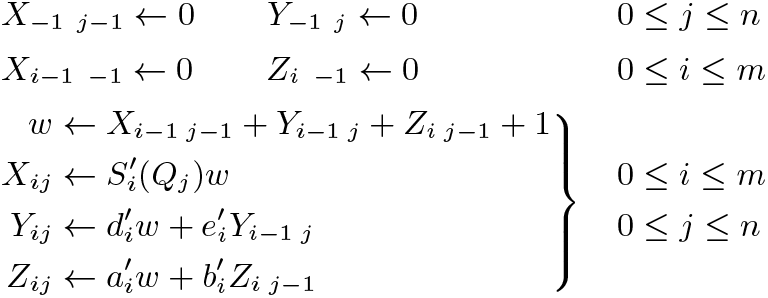

#### Algorithm 4

Sum of alignment extension probabilities

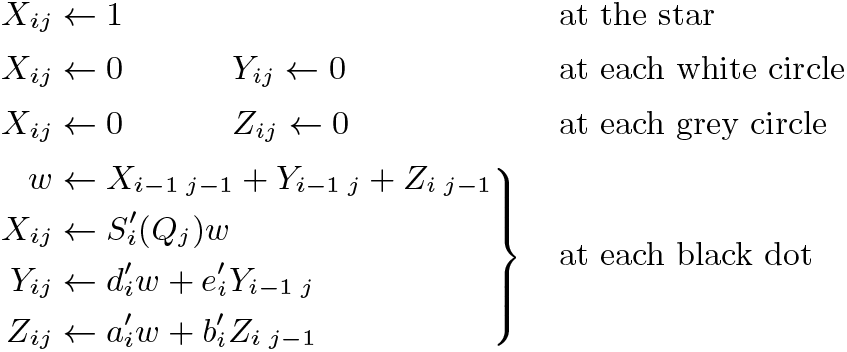

One alignment between two related regions does not hold all evidence that they are related. To judge whether they are related, it is better to calculate their probability without fixing an alignment (Frith 2024). In other words, use a score like this:

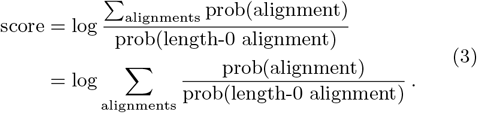

Which alignments should be included in the sum? The most extreme answer is to include all alignments between any parts of the sequence and profile. This is the most powerful way to judge if there are any related parts, but it does not tell us where those parts are, which is not very useful if the sequence or profile is long (for example, a chromosome).

Yu and Hwa (2001) suggested an “end-anchored” approach: include all alignments that start anywhere and end at some fixed coordinates (*i, j*) in the profile and the sequence. This enabled them to determine *E*-values. It is, however, unnatural: it would be appropriate if related regions have similarity with unclear starts but clear and sharp ends. A “mid-anchored” approach is more natural: include all alignments that pass through some fixed coordinates (*i, j*) (Frith 2024). This is approximated by a practical “heuristic mid-anchored” method: include all alignment extensions each side of an anchor, in a limited range (Fig. 2).

The algorithms for summing alignment probabilities depend on the precise definition of insertion and deletion probabilities. It is possible to define them in such a way that the algorithms have the same number of operations as Algorithms 1 and 2. This is likely the lowest number of operations any such affine-gap algorithm could have.

Algorithm 3 calculates the sum of probability ratios in Equation 3. It considers each length-*i* prefix of the profile and length-*j* prefix of the sequence:

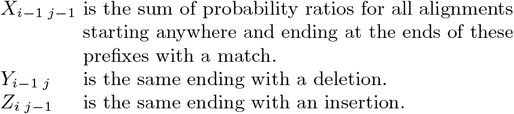

So, *w* is the sum of probability ratios of all alignments ending at the ends of these prefixes: the end-anchored probability ratio. The sum of all *w* values is the sum over all alignments. We can get mid-anchored probability ratios by combining end-anchored and start-anchored probability ratios (see the Supplement).

Algorithm 3 is a simple transformation of Algorithm 1, but it is not obvious that it validly sums alignment probabilities. To my knowledge, it has not been proposed before.

Algorithm 4 sums the probability ratios of all alignment extensions from an anchor: it is similar to Algorithm 2. There are alternative fast heuristics that do not extend from an anchor (Roddy et al. 2024): they can also use Algorithm 3 or 4 in a limited range, similar to the grey shapes in Fig. 2.

### Mitigating numeric overflow

Algorithm 3 may produce values larger than the computer can represent. This only happens when there is a strong similarity. Such overflow can be delayed by replacing the + 1 in Algorithm 3 with (for example) + 2^−63^. This simply makes all values calculated by the algorithm be multiplied by 2^−63^.

### Algorithm parameters in terms of probabilities

The algorithms are based on these letter and gap probabilities:

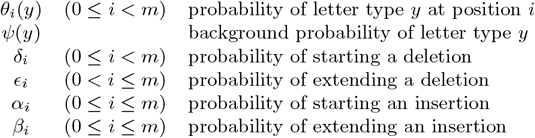

Surprisingly, *ϵ*_m_ is included: it implies deletion of the nonexisting position after the end of the profile. We will use it in the form 1 −*ϵ*_m_, the probability of not extending past the end.

These probabilities are defined precisely in the next section, but in the end we get these parameters for Algorithms 3 and 4:

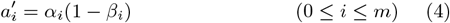

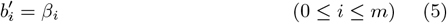

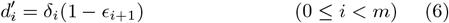

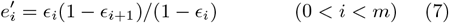

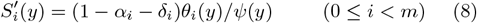

Finally, the score parameters for Algorithms 1 and 2 are

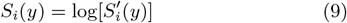

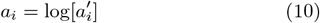

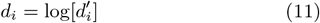

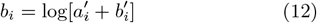

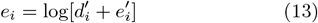

Equations 12 and 13 need explanation. As an example, consider two adjacent inserted letters. The algorithms allow them to come from either one length-2 insertion, or two length-1 insertions. We should sum these probabilities, to include as much evidence as possible. This is done by Algorithms 3 and 4, and also by Equations 12 and 13.

Algorithms 1–4 lack one feature of HMMER: it allows position-specific probabilities of letter types in insertions. Our algorithms can gain this feature with just one extra array lookup (see the Supplement).

### Precise definition of probabilities

Let us start by reviewing position-nonspecific alignment of two sequences (Frith 2020). Here, prob(alignment) means the product of probabilities of insertions, deletions, aligned letters, and unaligned letters in both sequences. A precise definition of these probabilities is shown in Fig. 3. Here, *α* is the insertion start probability, and *β* the insertion extend probability. Also, *δ* and *ϵ* are the deletion start and extend probabilities. There are also probabilities *τ* for ending the aligned region, *ω* for a flanking letter in the “query” sequence, and *η* for a flanking letter in the “reference” sequence. Overall, this defines probabilities for alignments between part of one sequence and part of the other.

**Figure 3.**
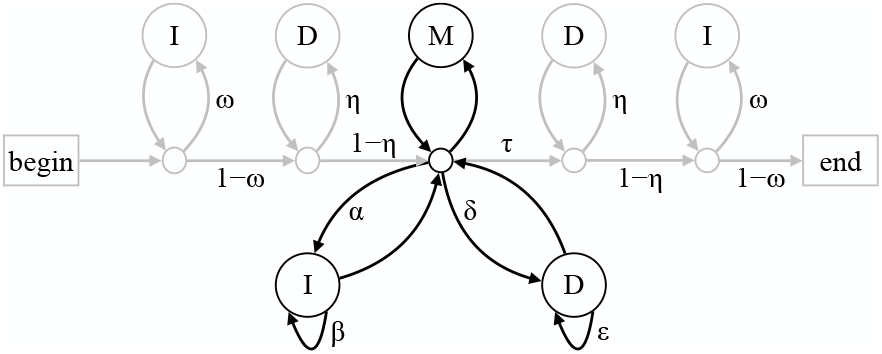
A scheme for assigning probability to: two sequences with one aligned region. The black part handles the aligned region; the grey parts handle unaligned flanks. Any path from **begin** to **end** represents an alignment: its probability is the product of the arrow probabilities (for example: *ω, η*) and letter probabilities. Each pass through a **D** matches the next letter in the “reference” sequence, with probability *ϕ*(*z*) for letter type *z*. Each pass through an **I** matches the next letter in the “query” sequence, with probability *ψ*(*z*) for letter type *z*. Each pass through **M** matches a reference letter *x* and a query letter *y*, with probability *π*(*x, y*).

Note that we apply these probabilities to two given sequences, of length *m* and *n*, and only consider paths that correspond to sequences of lengths *m* and *n*.

A definition of position-specific probabilities is shown in Fig. 4. It is derived by “unrolling” Fig. 3 for each position in the reference sequence, which is now the profile. An unusual feature is that we have arrows leading to empty space: these correspond to profile length ≠ *m*. As above, we only consider paths that correspond to profile length *m*, so we never use these arrows. It is possible to omit these arrows, but doing so makes the probabilities complicated (see the Supplement).

**Figure 4.**
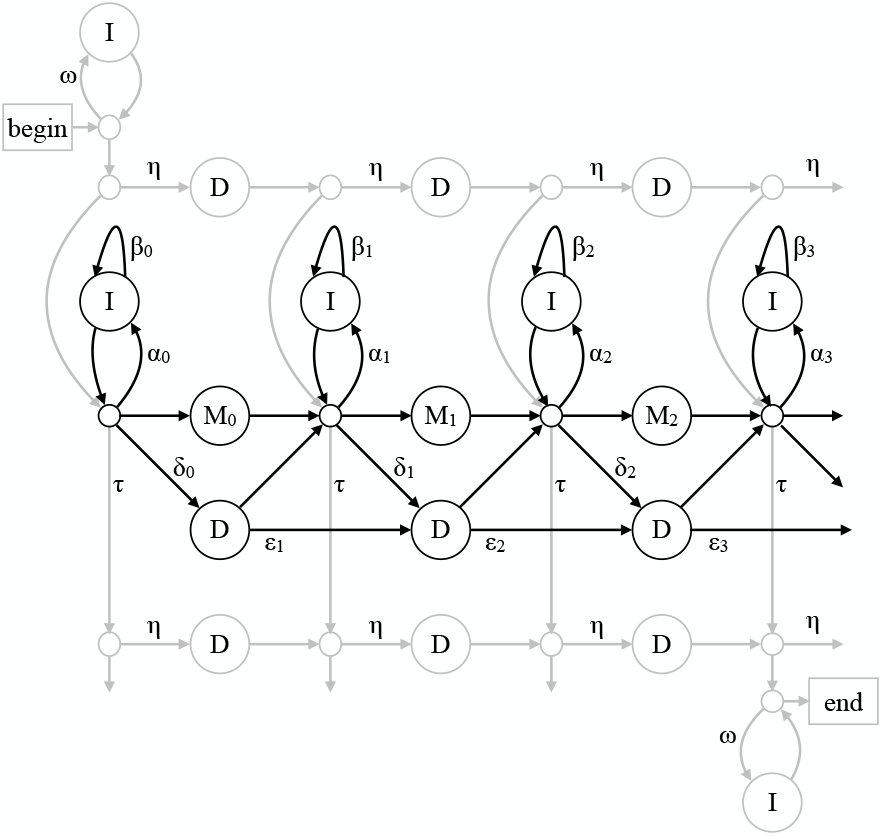
A scheme for assigning probability to: a sequence with one segment aligned to a profile. The black part handles alignment to the profile; the grey parts handle unaligned flanks. In this example, the profile has length *m* = 3. Any path from **begin** to **end** represents an alignment: its probability is the product of the arrow probabilities (for example: *ω, η*) and letter probabilities. Each pass through an **I** matches the next letter in the sequence, with probability *ψ*(*y*) for letter type *y*. Passing through **M**_*i*_ matches the next letter in the sequence, with probability *θ*_*i*_(*y*).

### Algorithm parameters from probabilities

We wish to calculate scores by Equation 2 or 3. Firstly, a length-0 alignment between the sequence and profile only traverses grey arrows in Fig. 4, never black arrows, so prob(length-0 alignment)

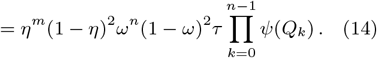

Next, consider a length-1 alignment that matches *Q*_j_ to *M*_i_:

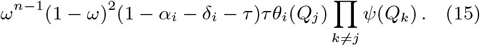

From dividing Equation 15 by Equation 14, we can infer that

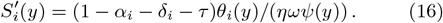

By similar reasoning, we can infer that

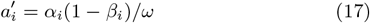

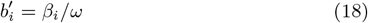

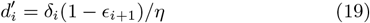

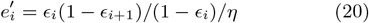

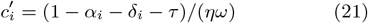

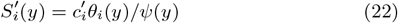

These are the same as Equations 4–8, assuming that *τ ≈* 0 and *ω ≈* 1 *≈ η*. These choices for *ω, η*, and *τ* mean there is no alignment length bias.

### Alignment length bias

The arrow probabilities may be biased towards long or short alignments, between a sequence and profile of given lengths. If *ω, η*, and *τ* are all high (almost 1), shorter alignments (with longer unaligned flanks) have higher probability. Conversely, if *ω, η*, and *τ* are all near 0, longer alignments (shorter flanks) have higher probability. There is no length bias if *ω ≈* 1 *≈ η* and *τ ≈* 0 (Frith 2020).

Lack of such bias may be desirable, for two reasons. Firstly, it means alignment is based purely on similarity, whereas long-bias aggressively aligns slightly dissimilar regions, and short-bias declines to align slightly similar regions. Secondly, it enables a way to determine *E*-values (Yu and Hwa 2001).

For gapless alignment (*α*_i_ = *δ*_i_ = 0), there is no alignment length bias when

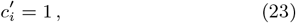

or equivalently

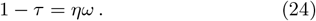

This means alignments of any length have the same product of arrow probabilities. Equation 23 is equivalent to Equation 1 in the work of Karlin and Altschul (1990). Also, *ω, η*, and *τ* are partly arbitrary: their values make no difference to our algorithms as long as they satisfy Equation 24.

As a point of comparison, HMMER3 is biased to longer alignments, at least in its gapless unihit mode. It has a probability *p <* 1 equivalent to our *ω*, without a counterbalancing equivalent of *τ >* 0 (Eddy 2008).

### Chance similarities in random sequences

Suppose we get a similarity score *s* by Equation 3. We would like to know the probability of getting a score *≥ s* by chance, in a random sequence. This can be estimated for a sequence of random independent letters, if:

- We use end-, start-, or mid-anchored scores.
- There is no alignment length bias.
- The random sequence has letter frequencies *ψ*(*y*). Based on the conjecture of Yu and Hwa, the *E*-value is:

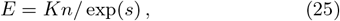

where *n* is the sequence length, in the limit where sequence and profile are long. Here, exp is defined to be the inverse of log (i.e. it depends on the base of the log). *K* is an unknown parameter, which is different for different profiles. *K* is roughly proportional to profile length, because a longer profile has more chances to match a random sequence, provided that the profile is not repetitive.

Equation 25 is expected to be less accurate for shorter sequences or profiles. One reason is an edge effect. The sum of probabilities of alignments anchored near the edge of the profile or sequence tends to be lower, because the alignments are limited by hitting the edge.

### Estimating *K*

To estimate *K* for a profile, we can generate random sequences with letter frequencies *ψ*(*y*), find the maximum score (over all possible (*i, j*) anchors) between the profile and each sequence, and fit *K* to these scores. The fitting can be done by maximum likelihood, which leads to this estimate of *K*:

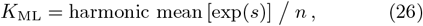

or by the method of moments:

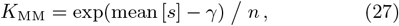

where *γ* is the Euler–Mascheroni constant (Yu et al. 2002). Longer random sequences make the estimates more accurate, but increase the computational cost. We can get more accuracy for less cost by adding “borders” to the random sequences, to suppress the edge effect. Altschul et al. (2001) suggested adding random letters to both ends of each sequence. This makes it necessary to ignore similarities that originate in the borders, else we trivially have longer random sequences.

A new idea here is to use a “wrapped border”. For example, if the random sequence length is 500 and the border length is 100, a copy of the sequence’s first 100 letters is appended to the sequence end (total length 600). This makes it unnecessary to ignore similarities originating in borders.

### HMMER3 *E*-value conjectures

The HMMER3 *E*-value conjectures seem to contradict the solution for gapless local alignment of two sequences (Karlin and Altschul 1990). For gapless alignment, we have scores *S*_xy_ for matching letter type *x* in one sequence and *y* in the other. If they are based on the probabilities in Fig. 3 (with *α* = *δ* = 0),

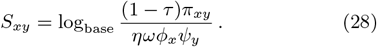

Other ways of defining local alignment probabilities would produce similar equations. If we compare two random sequences with letter probabilities Φ_x_ and Ψ_y_, the maximum alignment score follows a Gumbel distribution with a parameter *λ*, which is the solution of 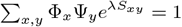 (Karlin and Altschul 1990). According to HMMER’s conjectures, *λ* = log[base]. But that is not always true.

### A prototype implementation

Prototype software named DUMMER (Dumb Uncomplicated Match ModelER) has been implemented (https://gitlab.com/mcfrith/seq-position-probs). It finds mid-anchored similarities without heuristics: it tries all possible (*i, j*) anchors (Algorithms 3 and S5). This makes it highly sensitive, and fast enough for some applications.

After finding a high-scoring anchor, DUMMER finds a representative alignment. It aligns letters whose probability of being aligned is *>* 0.5, among all alignments with that anchor (Miyazawa 1995). It finds these letters before and after the anchor. To find them after the anchor, it runs Algorithm 4 in a rectangle with top-left corner at the anchor (Fig. 2). The rectangle initially has side length 16: if this proves insufficient, it is doubled until sufficient. For each point (*i, j*) in the rectangle, Algorithm 4 calculates the sum of probabilities of alignments starting at the anchor and ending at that point. In finding anchors, DUMMER has already calculated the sum of probabilities of all alignments starting at each point. By combining these probabilities, we can get the probability that each pair of letters is aligned. When the sum of Algorithm 4’s probabilities reaches half the sum over all alignments starting at the anchor, there cannot be alignment probabilities *>* 0.5, so we can stop.

An early version found representative alignments for all high-scoring anchors (with *E*-value ≤ a threshold), then removed alignments that have any pair of aligned letters in common with a higher-scoring alignment. This proved too slow, because of many high-scoring anchors near each other. Thus, DUMMER only uses anchors whose score is maximum among anchors with the same *i* coordinate and *j* coordinate *±*31. This is still a bit slow.

Making a profile isn’t implemented yet, so DUMMER uses HMMER profiles (see the Supplement). HMMER defines insertion and deletion probabilities slightly differently than in Fig. 4, so they will be slightly suboptimal for our algorithms. One detail is that DUMMER’s background letter probabilities *ψ*(*y*) are set proportional to the geometric mean of *θ*_i_(*y*). This aims to stop the profile matching sequences that have no similarity other than similar letter abundances (Barrett et al. 1997).

## Results

### Chance similarities in random sequences

To find related regions between a sequence and a profile, we seek similarity scores higher than likely to occur by chance in a random sequence. We can estimate the probability of a similarity score occurring by chance, based on conjectures for infinite sequences and profiles. So we should test this for finite sequences and profiles.

This was tested by comparing some profiles to random sequences (Fig. 5). The DNA profiles (from Dfam) are of ancient transposons with relics in present-day genomes, and the protein profile (from Pfam) is for the Hobbit family of lipid transfer proteins. In most cases, the observed probability of getting a mid-anchored similarity score agrees closely with the expected probability (Fig. 5). This is also true for end- and start-anchored scores, and for other Dfam profiles (Supplementary Figs. S4–12).

**Figure 5.**
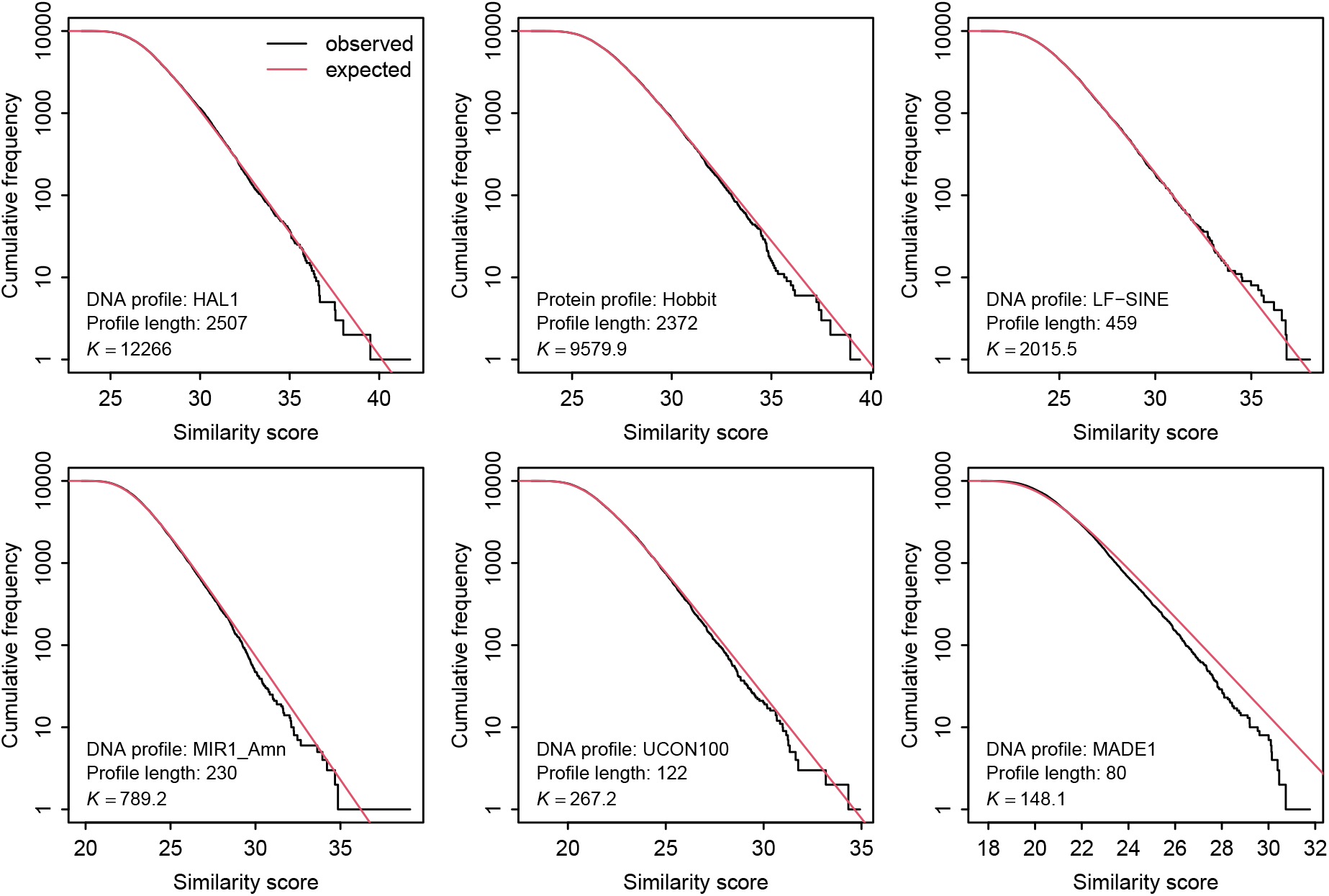
Distribution of similarity scores between between profiles and random sequences. For each profile, 10000 random sequences were generated, each of length 10000, using the profile’s background letter probabilities *ψ*(*y*). For each sequence, the maximum mid-anchored similarity score (over all possible (*i, j*) anchors) was found. The expected distribution depends on *K*, which was fitted to the observed similarity scores by the method of moments.

For the shortest profile (MADE1: 80 bases), the probability of getting a similarity score drops off with increasing score more steeply than expected (Fig. 5). This can be explained by edge effect: higher scores tend to involve longer alignments, which are increasingly limited by hitting the edge of the profile.

### A wrapped border improves *K* estimates

To get *E*-values, we need to estimate a parameter *K* for each profile, by finding similarity scores between the profile and random sequences. This estimation assumes that the random sequences are “long”, but it is not clear how long. To test this, *K* was estimated for a few profiles, with various sequence lengths (Fig. 6). The estimates of *K* decrease with decreasing sequence length. This can be explained by the edge effect: with shorter sequences, a greater fraction of potential anchors are near either edge of the sequence, and alignments with these anchors are limited by hitting the edge.

**Figure 6.**
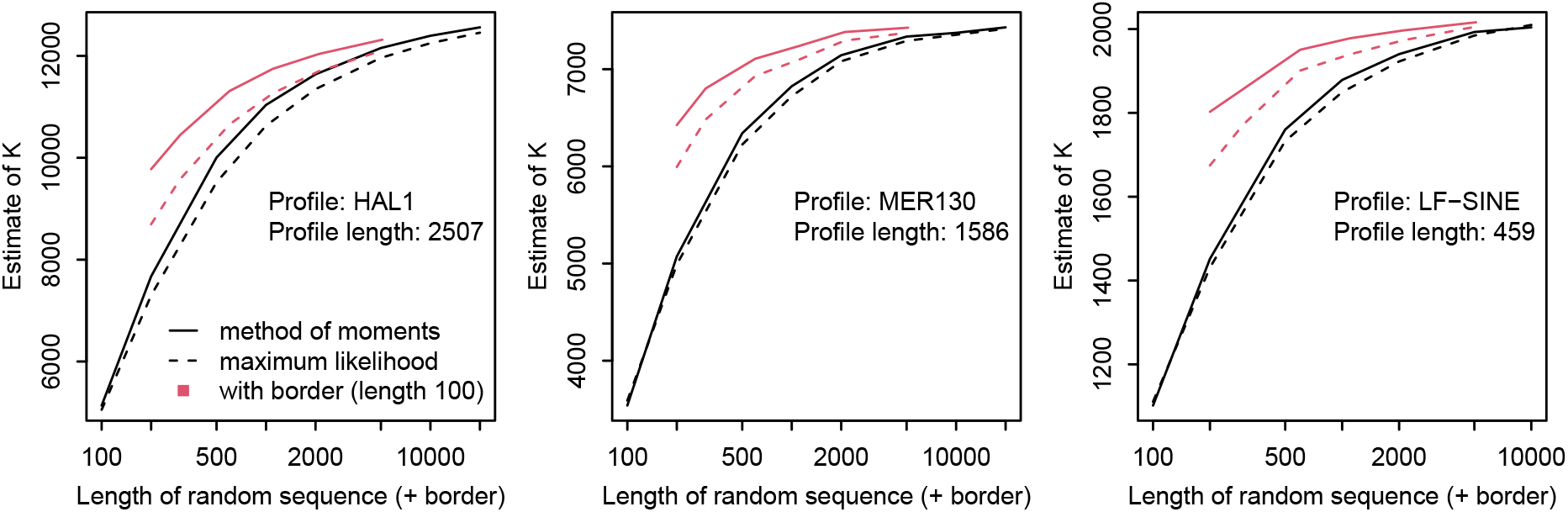
Value of *K* found by fitting to similarity scores between a profile and random DNA sequences. Different sequence lengths were tried (*x*-axis). For each length, the number of random sequences was 50000. Two fitting methods were tried: method of moments and maximum likelihood.

The edge effect can be suppressed by adding a “wrapped border” to each random sequence. Specifically, in these tests, a copy of each sequence’s first 100 letters was appended to the end of the sequence. This reduced the decrease in estimated *K* with decreasing sequence length (Fig. 6). Another, unexpected, result is that the method of moments works better than maximum likelihood, in these tests (Fig. 6).

### Sensitivity/specificity comparison to nhmmer

A first test of sensitivity was to detect relics of ancient HAL1 retrotransposons in human DNA. Some HAL1 sequences were extracted from the human genome (hg38), by taking the first 1000 segments (plus 100 nt flanks) annotated as HAL1 in `rmskOutCurrent` at the UCSC genome database. These annotations come from Dfam 2.0 and RepeatMasker v4.0.7 using nhmmer (nucleotide HMMER). So this test may be unfair in favor of nhmmer.

These HAL1 sequences were stored in one multi-fasta file, which was searched against the HAL1 profile. For each sequence, the highest mid-anchored score and its *E*-value were found, and the best nhmmer score and *E*-value were retained. These two *E*-values are comparable: they mean the expected number of matches in random sequences with the same length as *all* the HAL1 sequences. nhmmer was run with some nondefault options (`--F1 0.4 --F2 1 --F3 1 --nobias`), which turn off most of its heuristic shortcuts, to make it more sensitive but slow.

The mid-anchored *E*-values tend to be lower than the nhmmer *E*-values (Fig. 7 left). This means the mid-anchored approach can find the HAL1 matches with a lower rate of random matches (Fig. 5 left). So it is more sensitive, for any given rate of random matches (specificity).

**Figure 7.**
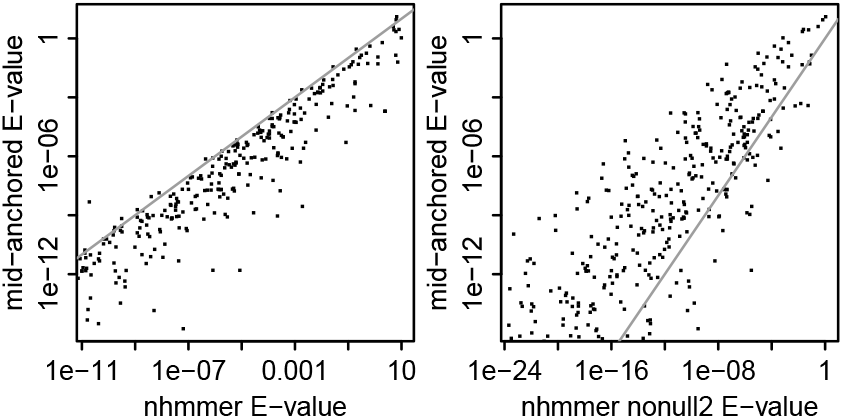
*E*-values for similarity between: the HAL1 profile, and human DNA with HAL1 RepeatMasker annotation. Each dot shows one DNA sequence. Each sequence has 3 kinds of *E*-value: mid-anchored, nhmmer, and nhmmer with option `--nonull2`.

As a further test, the nhmmer search was repeated with an extra option `--nonull2`, which turns off its correction for letter abundances. This time, the mid-anchored *E*-values tend to be higher than the nhmmer *E*-values (Fig. 7 right). So the “winner” depends on this nhmmer option: some other Dfam profiles produced similar results (Supplementary Figs. S13– 20).

Another test of sensitivity was to find parts of human chromosome 22 that are related to MIR1 Amn and LF-SINE (ancient retrotransposons). dummer was run with default options, and nhmmer with the same sensitive options as above. In this test, dummer found more matches than nhmmer to MIR1 Amn, for any given *E*-value threshold, and slightly more than nhmmer --nonull2 (Fig. 8). LF-SINE has just a handful of matches with *E*-value ≤ 0.01: dummer finds more than nhmmer (but less than nhmmer `--nonull2`) for a given *E*-value threshold.

**Figure 8.**
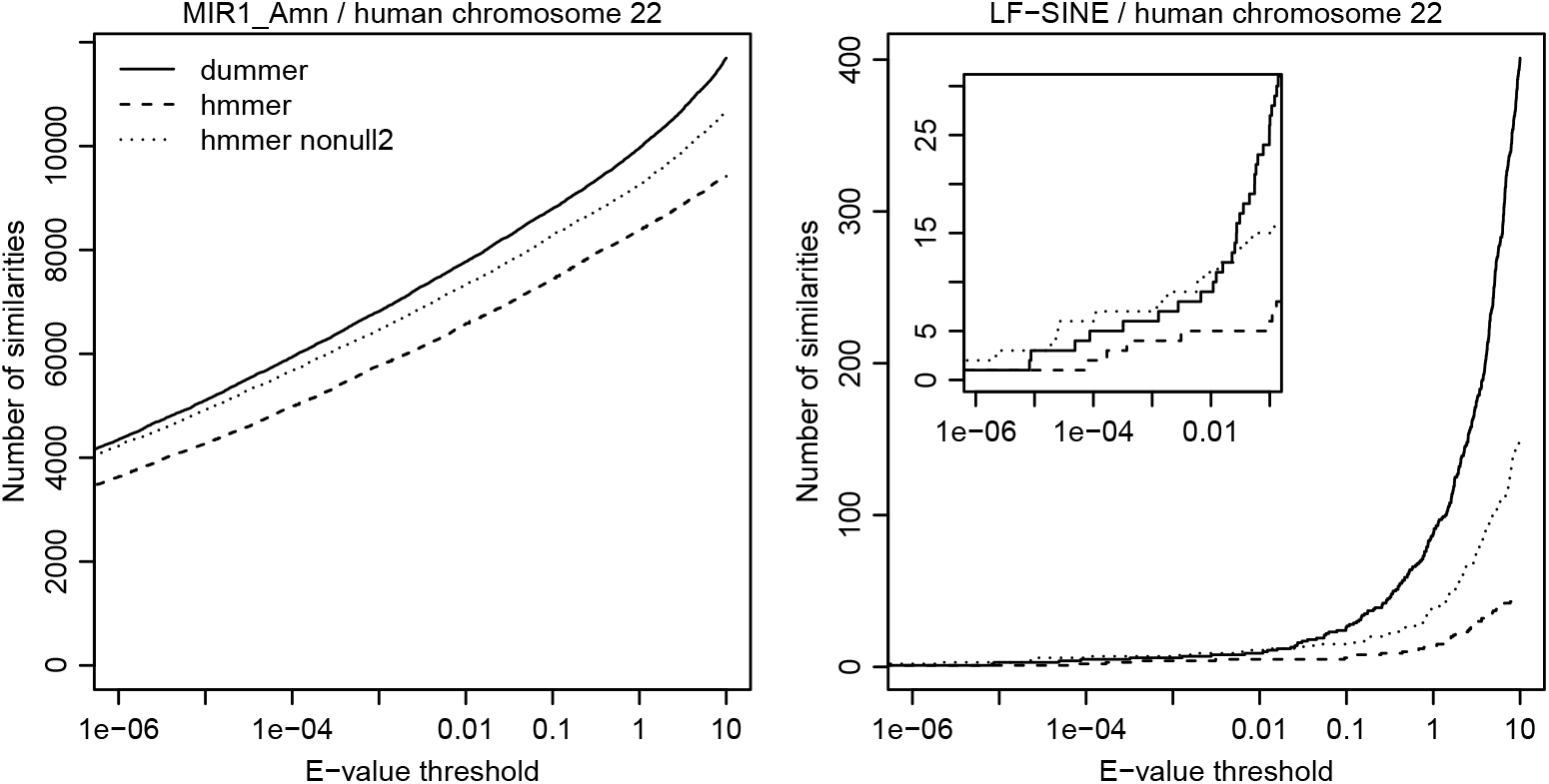
Number of matches to the MIR1 Amn profile (left) and the LF-SINE profile (right) in human chromosome 22, depending on *E*-value threshold. Three match-finding methods were tried: dummer, hmmer, and hmmer with option `--nonull2`. On the right, the inset shows the same plot zoomed in to the lower left.

Intriguingly, dummer revealed hundreds LF-SINE matches with borderline E-value ≤ 10 (Fig. 8). Some LF-SINE relics started performing various functions in common ancestors of all land vertebrates, and are ultraconserved (Bejerano et al. 2006). The dummer results indicate that the human genome has vastly more LF-SINE relics than previously known, at the statistical edge of detectability.

### Specificity in natural DNA

We have seen that dummer has high sensitivity for a given rate of matches to random DNA. But natural DNA does not consist of random independent letters with probabilities *ψ*(*y*). So we would like to know the rate of matches to unrelated natural DNA.

To test this, profiles of vertebrate transposons were searched against non-vertebrate DNA: chromosome 2L of the fruit fly *Drosophila melanogaster*, and chromosome 3 of the thale cress *Arabidopsis thaliana*. dummer found no hits with *E*-value *<* 0.1 for the LF-SINE and UCON100 profiles (Fig. 9), indicating good specificity. Not surprisingly, its hits do not exactly follow the *E*-values for random sequences: for example, 22 LF-SINE hits with *E*-value ≤ 10 to fruit fly DNA.

**Figure 9.**
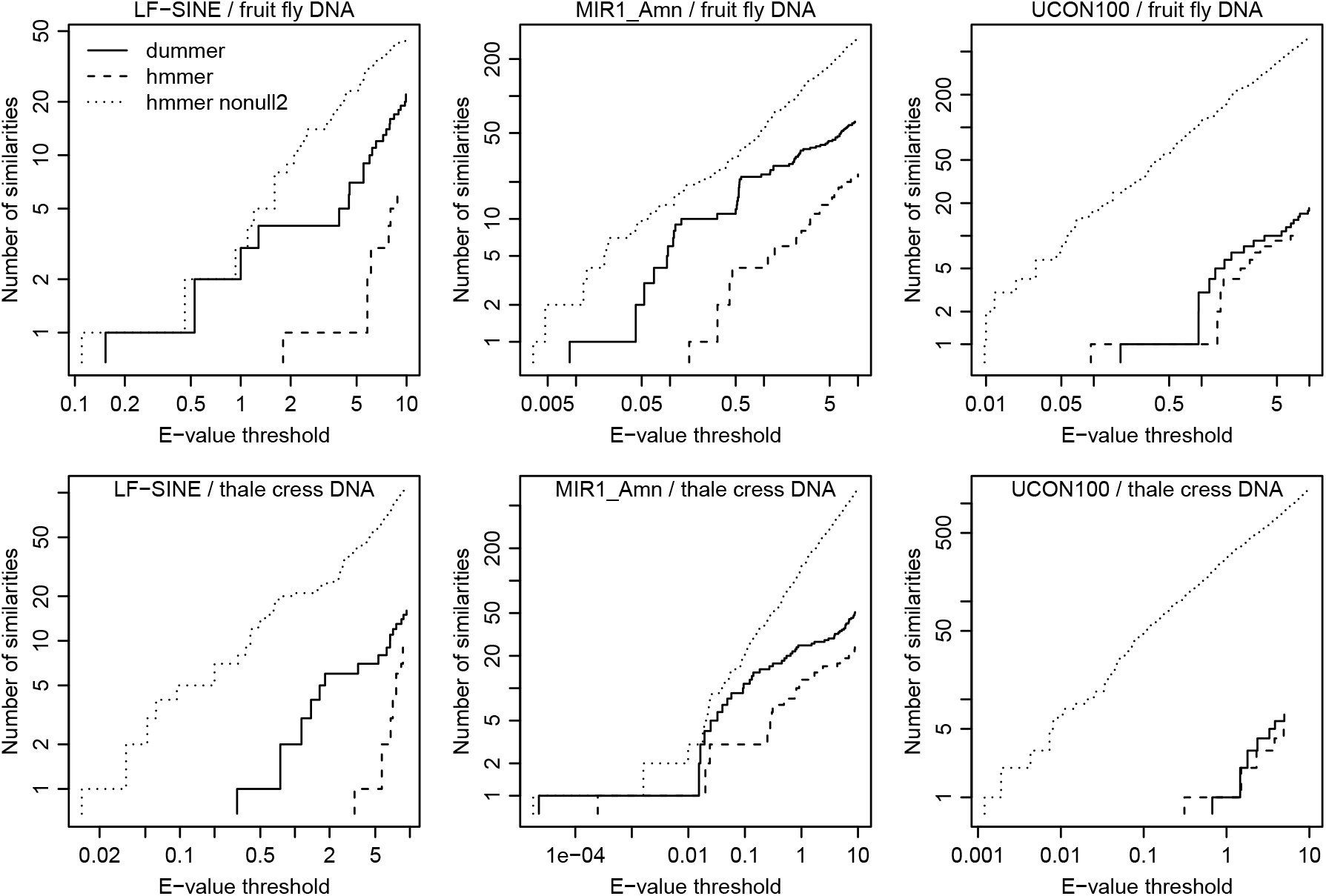
Number of matches between three profiles of vertebrate transposons and two non-vertebrate DNA sequences, depending on *E*-value threshold. Three match-finding methods were tried: dummer, hmmer, and hmmer with option `--nonull2`.

For the MIR1 Amn profile, however, dummer found matches with *E*-value 0.009 in fruit fly and 0.00002 in thale cress. These two hits (also nine of the top ten hits in fruit fly) overlap tRNA genes. MIR1 Amn is a SINE retroelement with a tRNA promoter, so these are true relationships.

### Computational cost

dummer was a few times slower than nhmmer (with slow options), and used vastly more memory (Table 1). So it cannot replace HMMER in general, but can be used for moderate-size searches. Its memory and run time are proportional to profile length *×* sequence length. HMMER uses heuristic shortcuts, and has received vastly more optimization effort. A widely useful tool requires heuristic shortcuts for adequate speed.

**Table 1.**
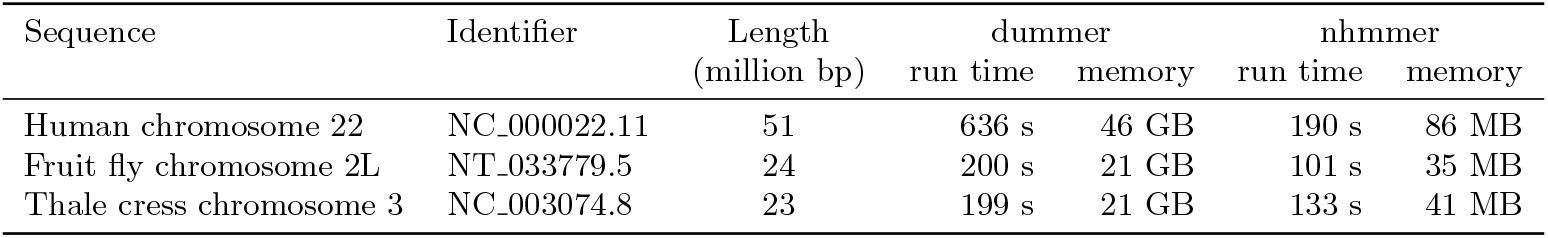
Time and memory for finding similarities between the MIR1 Amn profile and chromosomes.

## Discussion

This study shows how to achieve powerful HMMER-like detection of sequences related to profiles, without the complexity and minor quirks of HMMER. It is only necessary to implement a Gotoh-like algorithm (for example, Algorithm 3 or 4), whose parameters have simple formulas (Equations 4–8).

The DUMMER implementation has no heuristic shortcuts, making it sensitive and useful for moderate-size searches, and as a benchmark for faster heuristic software. The main value of this study will be if its simple methods are adpoted by widely-used heuristic tools.

DUMMER doesn’t avoid similarities of unrelated simple sequences, like atatatatatatatat, which evolve frequently and independently. This would be a problem for profiles with simple regions. The usual solution is to “mask” simple parts of sequences, though not all masking methods work equally well (Frith 2011).

One missing ingredient is a correction for *E*-value edge effect: the *E*-values are too conservative if the sequence or profile is short. Such corrections are possible, but seem a bit complicated (Yu and Hwa 2001, Eddy 2008).

This study neglects one important feature of HMMER: multihit comparison (Eddy 2008). We have assumed relationship between *one* part of a sequence and a profile, with unrelated flanks. This assumption can produce bad results when there are multiple related parts (Zhang et al. 1999). HMMER-style multihit comparison fixes the root cause of this problem, by assuming a *set* of related parts.

Finally, this study found evidence that the human genome has about 100 times more LF-SINE relics than was thought. In chromosome 22, 401 matches were found with *E*-values ≤ 10 (Fig. 8). About ten matches would be expected by chance in random DNA, and about 20 in unrelated natural DNA (Fig. 9). It would be interesting to detect these relics with greater confidence, perhaps by refining the LF-SINE profile, or by reverting evolution of the human genome sequence based on comparison with other mammals.

## Acknowledgments

I thank Travis Wheeler for explaining HMMER *E*-values, and Patrick Styll for investigating how to make a profile. This work was supported by JSPS KAKENHI grant number 22H04925 (PAGS).

## References

Altschul SF et al. (1997). Gapped BLAST and PSI-BLAST: a new generation of protein database search programs. Nucleic Acids Research 25:3389–3402.

Altschul SF et al. (2001). The estimation of statistical parameters for local alignment score distributions. Nucleic Acids Research 29:351–361.

Barrett C, Hughey R, Karplus K (1997). Scoring hidden Markov models. Bioinformatics 13:191–199.

Bejerano G et al. (2006). A distal enhancer and an ultraconserved exon are derived from a novel retroposon. Nature 441:87–90.

Cameron M, Williams HE, Cannane A (2004). Improved gapped alignment in BLAST. IEEE/ACM Transactions on Computational Biology and Bioinformatics 1:116–129.

Eddy SR (2008). A probabilistic model of local sequence alignment that simplifies statistical significance estimation. PLoS Computational Biology 4:e1000069.

Frith MC (2011). A new repeat-masking method enables specific detection of homologous sequences. Nucleic acids research 39:e23–e23.

Frith MC (2020). How sequence alignment scores correspond to probability models. Bioinformatics 36:408–415.

Frith MC (2024). A simple method for finding related sequences by adding probabilities of alternative alignments. Genome Research 34:1165–1173.

Frith MC, Ni S (2023). DNA conserved in diverse animals since the Precambrian controls genes for embryonic development. Molecular Biology and Evolution:msad275.

Gotoh O (1982). An improved algorithm for matching biological sequences. Journal of molecular biology 162:705– 708.

Gribskov M, McLachlan AD, Eisenberg D (1987). Profile analysis: detection of distantly related proteins. Proceedings of the National Academy of Sciences 84:4355–4358.

Karlin S, Altschul SF (1990). Methods for assessing the statistical significance of molecular sequence features by using general scoring schemes. Proceedings of the National Academy of Sciences 87:2264–2268.

Karplus K, Barrett C, Hughey R (1998). Hidden Markov models for detecting remote protein homologies. Bioinformatics 14:846–856.

Krogh A et al. (1994). Hidden Markov models in computational biology: Applications to protein modeling. Journal of molecular biology 235:1501–1531.

Liu D, Steinegger M (2023). Block Aligner: an adaptive SIMD-accelerated aligner for sequences and position-specific scoring matrices. Bioinformatics 39:btad487.

Miyazawa S (1995). A reliable sequence alignment method based on probabilities of residue correspondences. Protein Engineering, Design and Selection 8:999–1009.

Roddy JW, Rich DH, Wheeler TJ (2024). nail: software for high-speed, high-sensitivity protein sequence annotation. bioRxiv. doi: 10.1101/2024.01.27.577580.

Smith TF, Waterman MS (1981). Identification of common molecular subsequences. Journal of molecular biology 147:195–197.

Steinegger M, Söding J (2017). MMseqs2 enables sensitive protein sequence searching for the analysis of massive data sets. Nature biotechnology 35:1026–1028.

Steinegger M et al. (2019). HH-suite3 for fast remote homology detection and deep protein annotation. BMC bioinformatics 20:1–15.

Suzuki H, Kasahara M (2017). Acceleration of nucleotide semi-global alignment with adaptive banded dynamic programming. bioRxiv. doi: 10.1101/130633.

Wheeler TJ, Eddy SR (2013). nhmmer: DNA homology search with profile HMMs. Bioinformatics 29:2487–2489.

Yu YK, Bundschuh R, Hwa T (2002). Hybrid alignment: high-performance with universal statistics. Bioinformatics 18:864–872.

Yu YK, Hwa T (2001). Statistical significance of probabilistic sequence alignment and related local hidden Markov models. Journal of Computational Biology 8:249–282.

Zhang Z, Berman P, Miller W (1998). Alignments without low-scoring regions. Journal of Computational Biology 5:197– 210.

Zhang Z et al. (1999). Post-processing long pairwise alignments. Bioinformatics 15:1012–1019.

