## Supplement for "Simple and thorough detection of related sequences with position-varying probabilities of substitutions, insertions, and deletions"

Martin C. Frith

### Tracing back a maximum-score alignment

This section shows how to find an alignment extension with maximum score, after the score has been found by Algorithm 2. Let $(i_0, j_0)$ be the coordinates of the top-left-most black point in Fig. 2. We start with $(i, j)$ being the coordinates of a point with maximum $w$. This algorithm traces back a maximum-score alignment:

state $\leftarrow$ MATCH

While $i + j > i_0 + j_0$ :
  $y \leftarrow Y_{i-1\ j}$
  $z \leftarrow Z_{i\ j-1}$
  If state = DEL :  $y \leftarrow y - (d_i - e_i)$
  If state = INS :  $z \leftarrow z - (a_i - b_i)$

  If $X_{i-1\ j-1} \geq \max(y, z)$ :
    state $\leftarrow$ MATCH
  Else if $y \geq z$ :
    state $\leftarrow$ DEL
  Else :
    state $\leftarrow$ INS

  If state $\neq$ INS :  $i \leftarrow i - 1$
  If state $\neq$ DEL :  $j \leftarrow j - 1$

This algorithm's speed is less critical, because it visits only a small fraction of the black dots (Fig. 2).

### Varying probabilities of inserted letter types

This section modifies the algorithms, to allow position-specific probabilities of letter types in insertions. This means we have an extra set of probabilities $\chi_i(y)$: the probability of letter type $y$ in an insertion at position $i$ (Fig. S1). They can be handled by modifying Algorithms 1 to 4 into Algorithms S1 to S4. The parameters in Algorithms S3 and S4 are defined in terms of the probabilities like this:

$$T'_i(y) = \alpha_i(1-\beta_i)\chi_i(y)/\psi(y) \qquad (0 \le i \le m) \qquad \text{(S1)}$$
$$f'_i = \beta_i/[\alpha_i(1-\beta_i)] \qquad (0 \le i \le m) \qquad \text{(S2)}$$
$$d'_i = \delta_i(1-\epsilon_{i+1}) \qquad (0 \le i < m) \qquad \text{(S3)}$$
$$e'_i = \epsilon_i(1-\epsilon_{i+1})/(1-\epsilon_i) \qquad (0 \le i < m) \qquad \text{(S4)}$$
$$S'_i(y) = (1-\alpha_i-\delta_i)\theta_i(y)/\psi(y) \qquad (0 \le i < m) \qquad \text{(S5)}$$

The parameters in Algorithms S1 and S2 are defined like this:

$$S_i(y) = \log[S'_i(y)] \qquad \text{(S6)}$$
$$f_i = \log[f'_i + 1] \qquad \text{(S7)}$$
$$d_i = \log[d'_i] \qquad \text{(S8)}$$
$$T_i(y) = \log[T'_i(y)] \qquad \text{(S9)}$$
$$e_i = \log[d'_i + e'_i] \qquad \text{(S10)}$$

Algorithms S1 to S4 are almost as fast and simple as Algorithms 1 to 4, with just one extra operation (an array lookup).

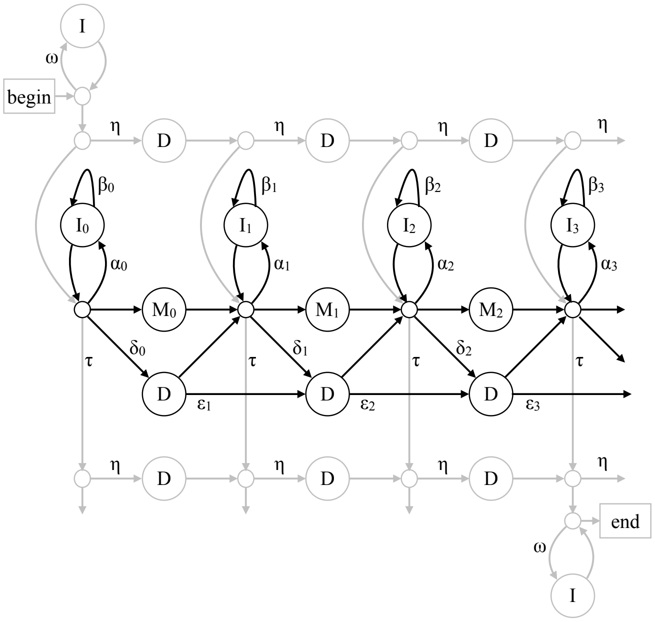


**Figure S1:** A scheme for assigning probability to: a sequence with one segment aligned to a profile. This is the same as Fig. 4, except it distinguishes insertions at different positions: $\mathbf{I}_0$, $\mathbf{I}_1$, ..., $\mathbf{I}_m$. Each pass through $\mathbf{I}_i$ matches the next letter in the sequence, with probability $\chi_i(y)$ for letter type $y$.

### Discussion

It is unclear if this modification is worthwhile. Krogh et al. (1993) found that it isn't. Eddy (1998) suggested it could be useful, if insertions have a general tendency, for example, to hydrophilic amino acids. On the other hand, Eddy (2011) mentioned that "Insert states emit residues with emission probabilities identical to a background distribution".

**Algorithm S1:** Maximum alignment score

$X_{-1\ j-1} \leftarrow -\infty \qquad Y_{-1\ j} \leftarrow -\infty \qquad 0 \le j \le n$

$X_{i-1\ -1} \leftarrow -\infty \qquad Z_{i\ -1} \leftarrow -\infty \qquad 0 \le i \le m$

$$\left.\begin{aligned} w &\leftarrow \max(X_{i-1\,j-1}, Y_{i-1\,j}, Z_{i\,j-1}, 0) \\ X_{ij} &\leftarrow w + S_i(Q_j) \\ Y_{ij} &\leftarrow \max(w + d_i, Y_{i-1\,j} + e_i) \\ Z_{ij} &\leftarrow \max(w, Z_{i\,j-1} + f_i) + T_i(Q_j) \end{aligned}\right\} \begin{aligned} 0 \le i \le m \\ 0 \le j \le n \end{aligned}$$

**Algorithm S2:** Maximum alignment extension score

$X_{ij} \leftarrow 0$ at the star

$X_{ij} \leftarrow -\infty \qquad Y_{ij} \leftarrow -\infty$ at each white circle

$X_{ij} \leftarrow -\infty \qquad Z_{ij} \leftarrow -\infty$ at each grey circle

$$\left.\begin{aligned} w &\leftarrow \max(X_{i-1\,j-1}, Y_{i-1\,j}, Z_{i\,j-1}) \\ X_{ij} &\leftarrow w + S_i(Q_j) \\ Y_{ij} &\leftarrow \max(w + d_i,\ Y_{i-1\,j} + e_i) \\ Z_{ij} &\leftarrow \max(w,\ Z_{i\,j-1} + f_i) + T_i(Q_j) \end{aligned}\right\} \text{at each black dot}$$

**Algorithm S3:** Sum of alignment probabilities

$X_{-1\ j-1} \leftarrow 0 \qquad Y_{-1\ j} \leftarrow 0 \qquad 0 \le j \le n$

$X_{i-1\ -1} \leftarrow 0 \qquad Z_{i\ -1} \leftarrow 0 \qquad 0 \le i \le m$

$$\left.\begin{aligned} w &\leftarrow X_{i-1\,j-1} + Y_{i-1\,j} + Z_{i\,j-1} + 1 \\ X_{ij} &\leftarrow S'_i(Q_j)w \\ Y_{ij} &\leftarrow d'_i w + e'_i Y_{i-1\,j} \\ Z_{ij} &\leftarrow (w + f'_i Z_{i\,j-1})T'_i(Q_j) \end{aligned}\right\} \begin{aligned} 0 \le i \le m \\ 0 \le j \le n \end{aligned}$$

**Algorithm S4:** Sum of alignment extension probabilities

$X_{ij} \leftarrow 1$ at the star

$X_{ij} \leftarrow 0 \qquad Y_{ij} \leftarrow 0$ at each white circle

$X_{ij} \leftarrow 0 \qquad Z_{ij} \leftarrow 0$ at each grey circle

$$\left.\begin{aligned} w &\leftarrow X_{i-1\,j-1} + Y_{i-1\,j} + Z_{i\,j-1} \\ X_{ij} &\leftarrow S'_i(Q_j)w \\ Y_{ij} &\leftarrow d'_i w + e'_i Y_{i-1\,j} \\ Z_{ij} &\leftarrow (w + f'_i Z_{i\,j-1})T'_i(Q_j) \end{aligned}\right\} \text{at each black dot}$$

### Sum of start-anchored alignment probabilities

Start-anchored is simply the opposite of end-anchored. Here, we sum the probabilities of all alignments that end anywhere and start at some fixed coordinates $(i, j)$ in the profile and the sequence. This can be done using Algorithm S5. It considers each suffix of the profile (with length $m-i$), and each suffix of the sequence (with length $n-j$). $\bar{W}_{ij}$ is the sum of probability ratios of all alignments starting at the starts of these suffixes.

**Algorithm S5:** Backward sum of alignment probabilities

$$\bar{W}_{m+1\ j+1} \leftarrow 0 \qquad \bar{Y}_{m+1\ j} \leftarrow 0 \qquad n \geq j \geq 0$$

$$\bar{W}_{i+1\ n+1} \leftarrow 0 \qquad \bar{Z}_{i\ n+1} \leftarrow 0 \qquad m \geq i \geq 0$$

$$\left.\begin{aligned} x &\leftarrow S'_i(Q_j)\bar{W}_{i+1\ j+1} \\ \bar{W}_{ij} &\leftarrow x + d'_i\bar{Y}_{i+1\ j} + a'_i\bar{Z}_{i\ j+1} + 1 \\ \bar{Y}_{ij} &\leftarrow \bar{W}_{ij} + e'_i\bar{Y}_{i+1\ j} \\ \bar{Z}_{ij} &\leftarrow \bar{W}_{ij} + b'_i\bar{Z}_{i\ j+1} \end{aligned}\right\} \quad \begin{aligned} m \geq i \geq 0 \\ n \geq j \geq 0 \end{aligned}$$

### Sum of mid-anchored alignment probabilities

The mid-anchored approach sums the probabilities of all alignments that pass through some fixed coordinates $(i, j)$ in the profile and the sequence. This can be done by first using Algorithm S5, and then using Algorithm 3 with one extra line shown in red:

$$\left.\begin{aligned} w &\leftarrow X_{i-1\ j-1} + Y_{i-1\ j} + Z_{i\ j-1} + 1 \\ \widetilde{W}_{ij} &\leftarrow w\bar{W}_{ij} \\ X_{ij} &\leftarrow S'_i(Q_j)w \\ Y_{ij} &\leftarrow d'_i w + e'_i Y_{i-1\ j} \\ Z_{ij} &\leftarrow a'_i w + b'_i Z_{i\ j-1} \end{aligned}\right\} \quad \begin{aligned} 0 \leq i \leq m \\ 0 \leq j \leq n \end{aligned}$$

Here, $\widetilde{W}_{ij}$ is the sum of probability ratios in Equation 3, over all alignments passing through $(i, j)$. So the score is $\log \widetilde{W}_{ij}$.

### Probabilities without unused arrows

Fig. S2 shows a precise definition of alignment probabilities, like Fig. 4, but without unused arrows. We require that all length-0 alignments (which only traverse grey arrows) have the same probability. This means that

$$(1-\eta_i)\tau_i = \eta_i(1-\eta_{i+1})\tau_{i+1} \qquad (0 \le i < m)\,. \tag{S11}$$

Here, $\eta_m$ is defined to be 0. So, we can calculate all the $\eta_i$ values from the $\tau_i$ values:

$$\eta_i = 1/[1+(1-\eta_{i+1})\tau_{i+1}/\tau_i] \qquad (0 \le i < m)\,. \tag{S12}$$

Also,

$$\text{prob(length-0 alignment)} = \omega^n(1-\omega)^2(1-\eta_0)\tau_0 \prod_{k=0}^{n-1}\psi(Q_k)\,. \tag{S13}$$

Next, similarly to the main text, we can determine the algorithm parameters in terms of the probabilities:

$$a'_i = \alpha_i(1-\beta_i)/\omega \qquad (0 \le i \le m) \tag{S14}$$
$$b'_i = \beta_i/\omega \qquad (0 \le i \le m) \tag{S15}$$
$$d'_i = \delta_i(1-\epsilon_{i+1})/(\tau_i/\tau_{i+1}) \qquad (0 \le i < m) \tag{S16}$$
$$e'_i = \epsilon_i(1-\epsilon_{i+1})/(1-\epsilon_i)/(\tau_i/\tau_{i+1}) \qquad (0 < i < m) \tag{S17}$$
$$c'_i = (1-\alpha_i-\delta_i-\tau_i)/(\omega\tau_i/\tau_{i+1}) \qquad (0 \le i < m) \tag{S18}$$
$$S'_i(y) = c'_i\theta_i(y)/\psi(y) \qquad (0 \le i < m) \tag{S19}$$

Here, $\epsilon_m$ is defined to be 0.

#### Position-nonspecific parameters

The problem with this scheme is that the probabilities are unintuitive. In particular, position-nonspecific parameters require position-varying probabilities. Position-nonspecific parameters mean that

$$a'_i = a' \qquad b'_i = b' \qquad d'_i = d' \qquad e'_i = e' \qquad c'_i = c'\,.$$

It follows that $\beta_i$ and $\alpha_i$ are non-varying:

$$\beta_i = \beta = \omega b'\,, \tag{S20}$$
$$\alpha_i = \alpha = \omega a'/(1-\omega b')\,. \tag{S21}$$

The other arrow probabilities, however, do vary:

$$\delta_i = \frac{d'(1-\alpha)}{d'+(\omega c'+\tau_{i+1})(1-\epsilon_{i+1})} \tag{S22}$$
$$\tau_i = \frac{(1-\alpha)\tau_{i+1}(1-\epsilon_{i+1})}{d'+(\omega c'+\tau_{i+1})(1-\epsilon_{i+1})} \tag{S23}$$
$$\epsilon_i = \frac{e'(1-\alpha)}{d'+(\omega c'+\tau_{i+1})(1-\epsilon_{i+1})+e'(1-\alpha)} \tag{S24}$$

These equations let us calculate $\delta_i$, $\tau_i$, and $\epsilon_i$ in terms of $\tau_{i+1}$ and $\epsilon_{i+1}$. Also, we know that $\tau_m = 1-\alpha$ and $\epsilon_m = 0$.

Fig. S3 shows an example, for some reasonable values of $a'$, $b'$, $d'$, $e'$, and $c'$. We can see that the values of $\delta_i$, $\tau_i$, and $\epsilon_i$ depend also on $\omega$. Here too, the probabilities are partly arbitrary: their values make no difference to our algorithms as long as the left-hand sides of Equations S14–S19 don't change.

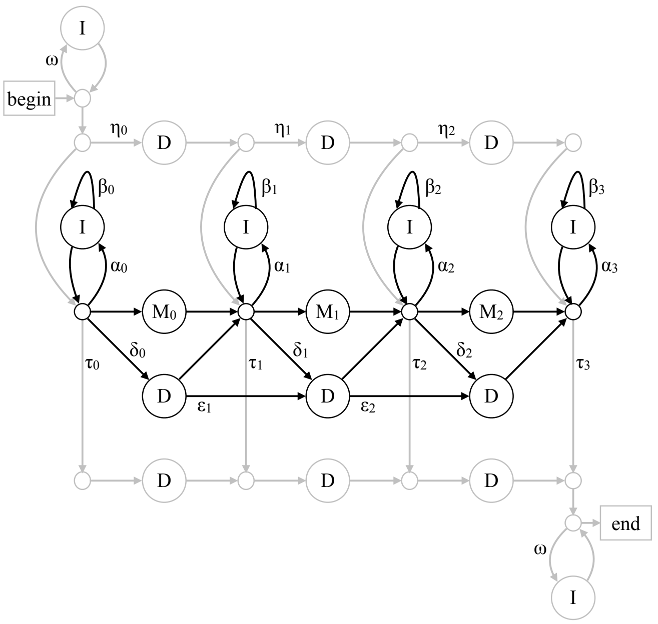


**Figure S2:** A scheme for assigning probability to: a sequence with one segment aligned to a profile. The black part handles alignment to the profile; the grey part handles unaligned flanks. In this example, the profile has length $m = 3$. Any path from **begin** to **end** represents an alignment: its probability is the product of the arrow probabilities (for example: $\omega$, $\eta_0$) and letter probabilities. Each pass through an **I** matches the next letter in the sequence, with probability $\psi(y)$ for letter type $y$. Passing through $\mathbf{M}_i$ matches the next letter, with probability $\theta_i(y)$.

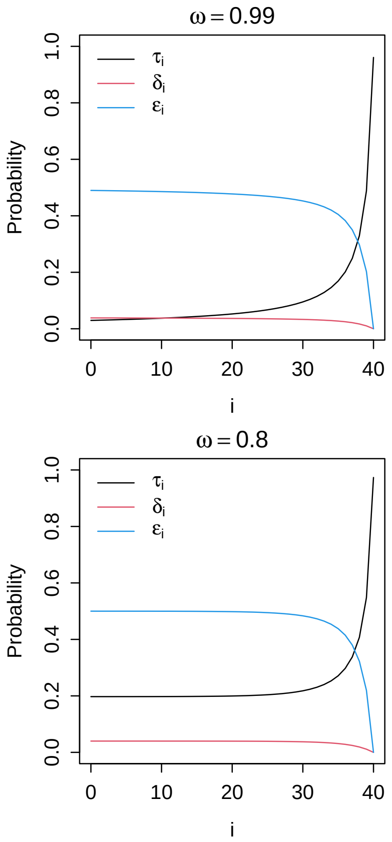


**Figure S3:** Values of $\tau_i$, $\delta_i$, and $\epsilon_i$, when $b' = e' = 0.5$, $a' = d' = 0.02$, and $c' = 0.92$, in a profile of length $m = 40$.

### Making a profile of some sequences

We would like to make a profile of some sequences, for example, the sequences in Fig. 1. The standard method is the Baum-Welch algorithm, which tries to find profile probabilities that maximize the probability of the sequences (Krogh et al. 1994). It finds a local maximum, but not necessarily a global maximum. Here is the basic Baum-Welch algorithm:

1. We need an initial guess for the profile, with some length $m$, and initial values for all the probabilities.
2. For each sequence, get the expected number of times to traverse each arrow, and the expected number of times to see each letter type in each position.
3. Add each type of expected count over all sequences.
4. Make new estimates for all the profile probabilities, which maximize the likelihood of the expected counts.
5. Go to step 2, and repeat until the profile stops changing.

This basic algorithm has the following problems and countermeasures.

- Some of the sequences may be closely related, for example, we may have mammal sequences, including several from apes that are 96% identical. The profile will be biased towards these similar sequences. A popular countermeasure is "position-based sequence weights", which down-weights similar sequences (S Henikoff and JG Henikoff 1994). It seems to require an alignment of the sequences: it would be convenient to require only the expected counts.
- The profile may get "over-fitted" to the sequences. In particular, if the sequences lack a letter type at a position, its probability will be zero, which seems too harsh. The solution is to use prior probabilities for the letter and arrow probabilities (Krogh et al. 1993). One method is "maximum a posteriori" (MAP) estimation, which seems a reasonable generalization of maximizing the probability of the sequences. Another method is "posterior mean" estimation, whose rationale seems less clear to me.
- We do not actually want the profile to fit the sequences. The aim is to find more-distantly related sequences. This can be addressed by "entropy weighting", which controls the weight given to the sequences versus the prior probabilities (Karplus et al. 1998).
- We may get stuck in a local optimum. One countermeasure is to make a good initial guess, for example, based on an alignment such as Fig. 1. Another countermeasure is to add random noise (like simulated annealing) to escape local optima.
- The initial profile length $m$ may not be ideal. This can be addressed by "model surgery": remove profile positions that are usually deleted, and add positions where there are usually insertions (Krogh et al. 1994). This may also help to avoid local optima.
- We may not get the most useful values for the background letter probabilities, $\psi(y)$. Barrett et al. (1997) suggested setting $\psi(y)$ proportional to the geometric mean of $\theta_i(y)$. This aims to stop the profile matching sequences that have no similarity other than similar letter abundances.

### Calculating the expected counts

Let us see how to do step 2 of the algorithm, which is unchanged by most of the countermeasures. This step gets $E[\heartsuit]$ (for example, $E[\beta_i]$): the expected count of the event with probability labeled $\heartsuit$. Let's define labels for unlabeled arrows in Fig. 4:

$$\begin{aligned}\widehat{\beta}_i &= 1 - \beta_i\,, \\ \widehat{\epsilon}_i &= 1 - \epsilon_i\,, \\ \gamma_i &= 1 - \alpha_i - \delta_i - \tau_i\,.\end{aligned}$$

We can get each $E[\heartsuit]$ from the "forward" Algorithm 3 and the "backward" Algorithm S5. Here, $v$ is defined to be the sum of all the $w$ values calculated by Algorithm 3.

$$E[\widehat{\beta}_i] = \left(\sum_{j=0}^{n} Z_{i\;j-1}\bar{W}_{ij}\right)/v \qquad (0 \le i \le m) \quad \text{(S25)}$$

$$E[\beta_i] = \left(\sum_{j=0}^{n} Z_{i\;j-1}\bar{Z}_{ij}\right)/v - E[\widehat{\beta}_i] \qquad (0 \le i \le m) \quad \text{(S26)}$$

$$E[\alpha_i] = E[\widehat{\beta}_i] \qquad (0 \le i \le m) \quad \text{(S27)}$$

$$E[\widehat{\epsilon}_i] = \left(\sum_{j=0}^{n} Y_{i-1\;j}\bar{W}_{ij}\right)/v \qquad (0 \le i \le m) \quad \text{(S28)}$$

$$E[\epsilon_i] = \left(\sum_{j=0}^{n} Y_{i-1\;j}\bar{Y}_{ij}\right)/v - E[\widehat{\epsilon}_i] \qquad (0 \le i \le m) \quad \text{(S29)}$$

$$E[\delta_i] = E[\widehat{\epsilon}_{i+1}] + E[\epsilon_{i+1}] - E[\epsilon_i] \qquad (0 \le i < m) \quad \text{(S30)}$$

$$E[\theta_i(y)] = \left(\sum_{j|Q_j=y} X_{ij}\bar{W}_{i+1\;j+1}\right)/v \qquad (0 \le i < m) \quad \text{(S31)}$$

$$E[\gamma_i] = \sum_{y} E[\theta_i(y)] \qquad (0 \le i < m) \quad \text{(S32)}$$

These edge values are always 0: $E[\widehat{\epsilon}_0]$, $E[\epsilon_0]$, and $E[\epsilon_m]$.

In the end, we will not get good values for $\widehat{\epsilon}_m$ or $\alpha_m$. They will be high (1 without prior probabilities). For example, high $\widehat{\epsilon}_m$ causes a bias in favor of deletions ending at the end of the profile. So perhaps $\widehat{\epsilon}_m$ and $\alpha_m$ should be determined by prior probabilities alone. I am also unsure if we will get useful values for $\alpha_0$, $\beta_0$, and $\beta_m$.

### Probabilities from HMMER profile files

We can read position-specific probabilities from a HMMER profile file. HMMER defines insertion and deletion probabilities differently than in Fig. 4, so they likely aren't optimal for this study's algorithms.

For a length-$m$ profile, a HMMER3/f file specifies these probabilities, for $0 \le k \le m$:

| | |
|---|---|
| $M_k \to I_k$ | probability of starting an insertion |
| $I_k \to I_k$ | probability of extending an insertion |
| $M_k \to D_{k+1}$ | probability of starting a deletion |
| $D_k \to D_{k+1}$ | probability of extending a deletion |

These ones are always zero: $D_0 \to D_1$, $M_m \to D_{m+1}$, and $D_m \to D_{m+1}$. The file also specifies $e_k(y)$, the probability of letter type $y$ at position $k$, for $0 < k \le m$.

The present study got probabilities from a HMMER file like this:

$$\alpha_k = (M_k \to I_k) \qquad 0 \le k \le m \qquad \text{(S33)}$$

$$\beta_k = (I_k \to I_k) \qquad 0 \le k \le m \qquad \text{(S34)}$$

$$\delta_k = (M_k \to D_{k+1}) \qquad 0 \le k < m \qquad \text{(S35)}$$

$$\epsilon_k = (D_k \to D_{k+1}) \qquad 0 < k < m \qquad \text{(S36)}$$

$$\theta_k(y) = e_{k+1}(y) \qquad 0 \le k < m \qquad \text{(S37)}$$

$$\epsilon_m = \underset{k=1}{\overset{m-1}{\text{geometric mean}}}[\epsilon_k] \qquad \text{(S38)}$$

$$\psi(y) \propto \underset{k=0}{\overset{m-1}{\text{geometric mean}}}[\theta_k(y)] \qquad \text{(S39)}$$

### Reproducing the results

The results can be reproduced as follows. The tested profiles and sequences are available at https://gitlab.com/mcfrith/seq-position-probs.

#### HMMER

For Fig. 7, nhmmer (from HMMER v3.4) was run like this:

```
nhmmer --watson --F1 0.4 --F2 1 --F3 1 --nobias
       --noali profile.hmm sequences.fasta
```

`--watson` makes it search forward DNA strands only.

`--noali` just makes the output more compact.

nhmmer was then run again, adding option `--nonull2`. For Figs. 8 and 9, nhmmer was run in the same ways, except that `--cpu 1` was used, and neither `--watson` nor `--noali` were used.

#### DUMMER

Fig. 5 can be reproduced with DUMMER v1.0.0 like this:

```
dummer -t10000 -l10000 -b0 profile.hmm
```

This specifies 10000 random sequences of length 10000, with no border. Fig. 6 can be reproduced similarly, by varying the sequence length `-l` and border length `-b`. Fig. 7 can be reproduced like this:

```
dummer -t10000 -l10000 -b0 -s1 -e0 profile.hmm
                                    sequences.fasta
```

`-s1` makes it search forward DNA strands only.

`-e0` makes it report the strongest similarity per sequence, instead of all similarities with $E$-value $\le$ a threshold.

Figs. 8 and 9 can be reproduced like this:

```
dummerl profile.hmm sequences.fasta
```

`dummerl` is the low-memory version of dummer: the only difference is that it uses single-precision instead of double-precision floating-point numbers.

### References


Barrett C, Hughey R, Karplus K (1997). Scoring hidden Markov models. *Bioinformatics* **13**:191–199.

Eddy SR (2011). Accelerated profile HMM searches. *PLoS computational biology* **7**:e1002195.

Eddy SR (1998). Profile hidden Markov models. *Bioinformatics* **14**:755–763.

Henikoff S, Henikoff JG (1994). Position-based sequence weights. *Journal of molecular biology* **243**:574–578.

Karplus K, Barrett C, Hughey R (1998). Hidden Markov models for detecting remote protein homologies. *Bioinformatics* **14**:846–856.

Krogh A et al. (1993). *Hidden Markov models in computational biology: Applications to protein modeling.* Tech. rep. University of California at Santa Cruz.

Krogh A et al. (1994). Hidden Markov models in computational biology: Applications to protein modeling. *Journal of molecular biology* **235**:1501–1531.

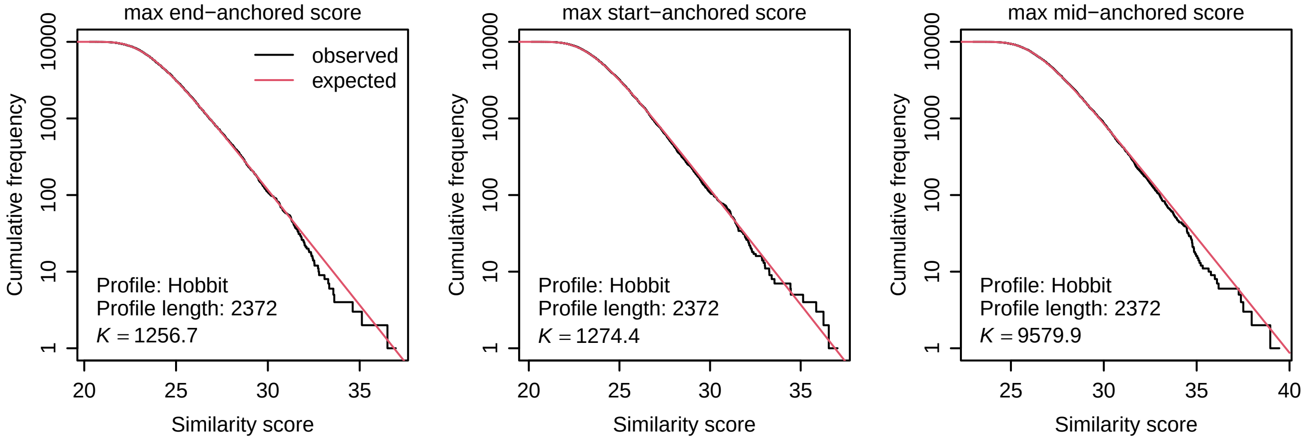


**Figure S4:** Distributions of similarity scores between the Hobbit profile (PF10344.16) and random protein sequences. 10000 random sequences were generated, each of length 10000, using the profile's background letter probabilities $\psi(y)$. For each sequence, the maximum end-anchored, start-anchored, and mid-anchored similarity score was found. The expected distribution depends on $K$, which was fitted to the observed similarity scores by the method of moments.

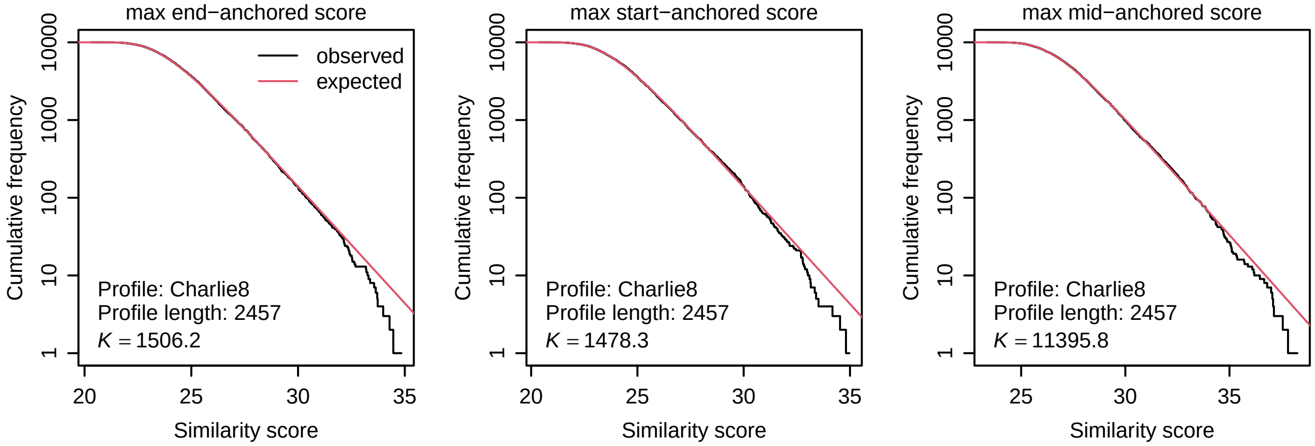


**Figure S5:** Distributions of similarity scores between the Charlie8 profile (DF000000107.4) and random DNA sequences. 10000 random sequences were generated, each of length 10000, using the profile's background letter probabilities $\psi(y)$. For each sequence, the maximum end-anchored, start-anchored, and mid-anchored similarity score was found. The expected distribution depends on $K$, which was fitted to the observed similarity scores by the method of moments.

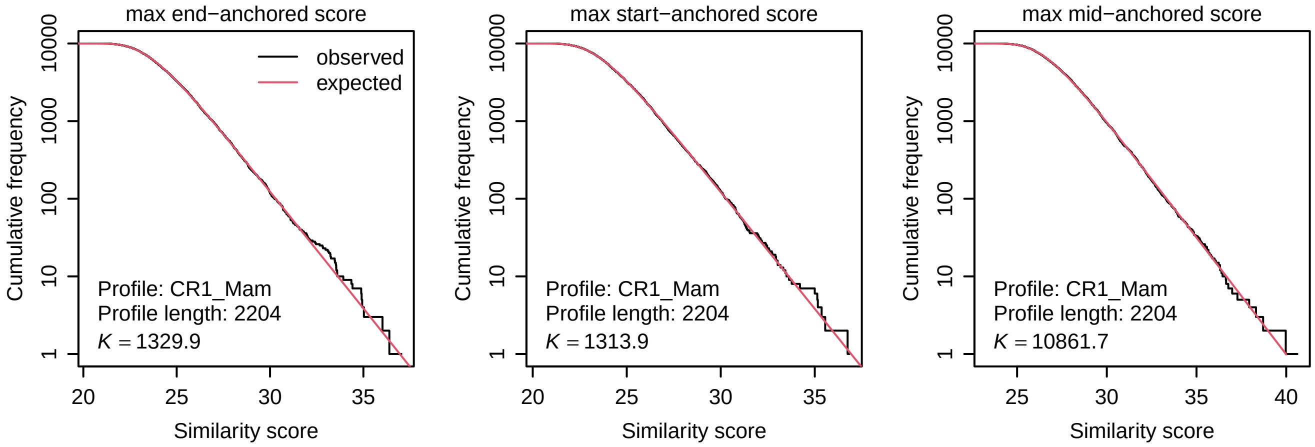


**Figure S6:** Distributions of similarity scores between CR1_Mam (DF000000110.4) and random DNA sequences. See the Fig. S5 legend.

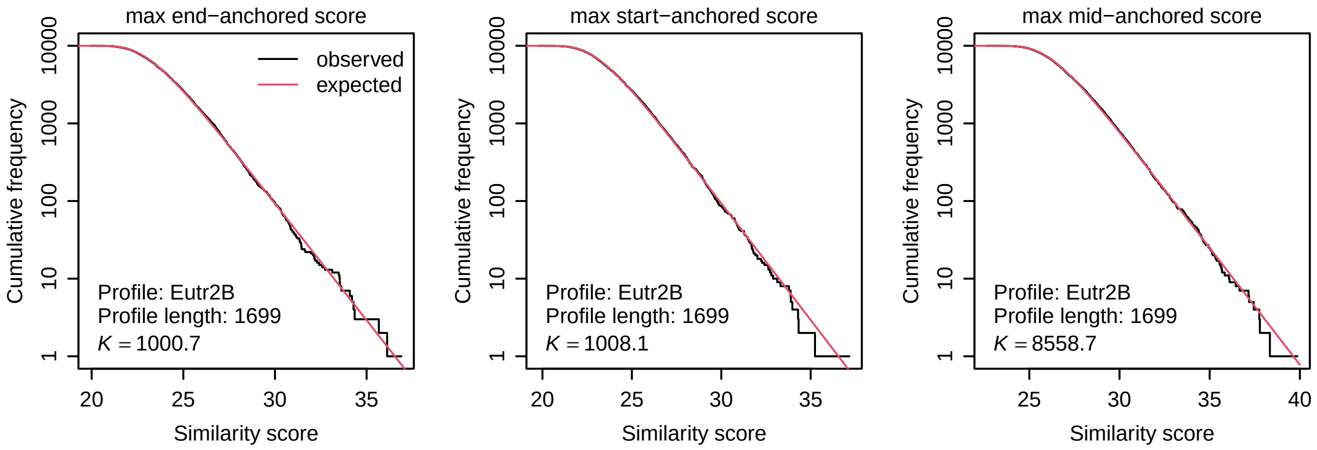


**Figure S7:** Distributions of similarity scores between Eutr2B (DF000001264.2) and random DNA sequences.

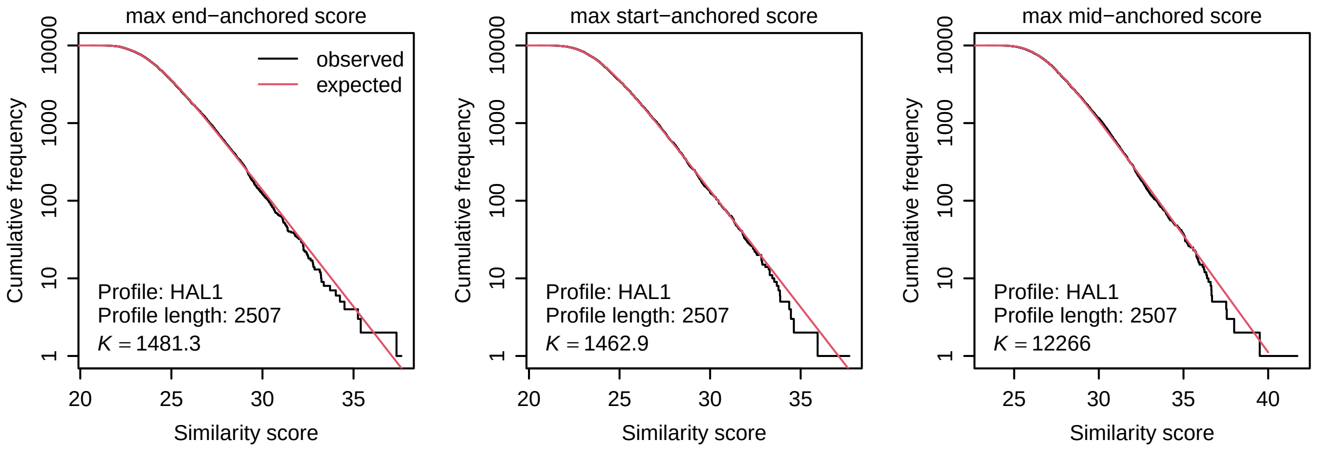


**Figure S8:** Distributions of similarity scores between HAL1 (DF000000154.4) and random DNA sequences.

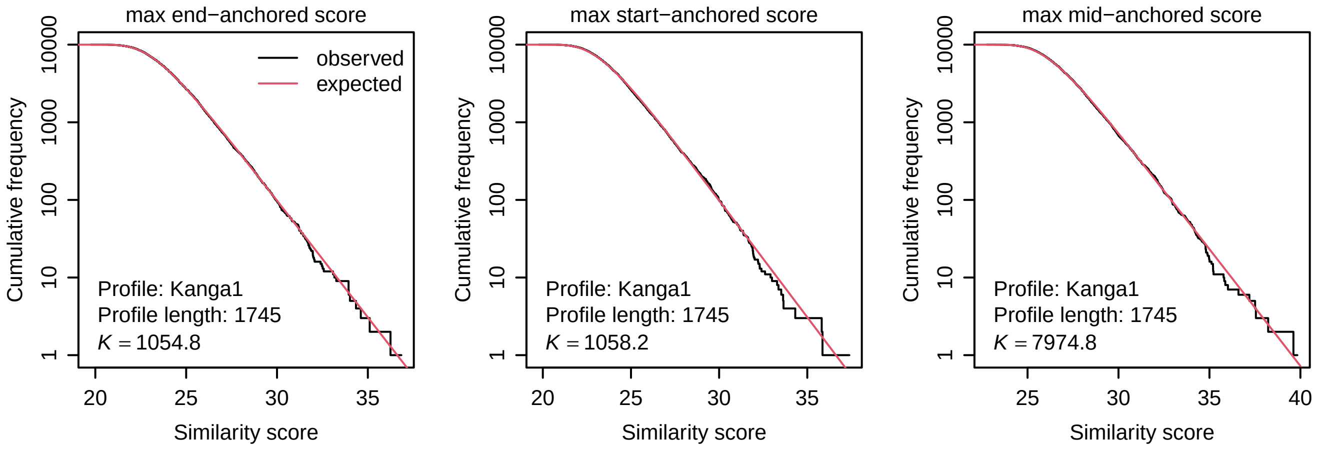


**Figure S9:** Distributions of similarity scores between Kanga1 (DF000000218.4) and random DNA sequences.

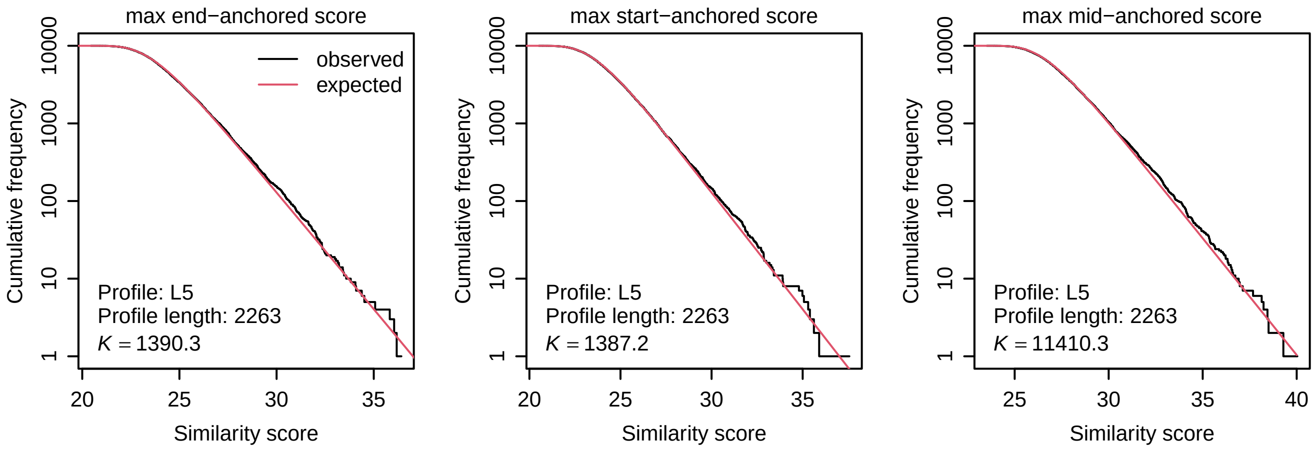


**Figure S10:** Distributions of similarity scores between L5 (DF000000367.5) and random DNA sequences.

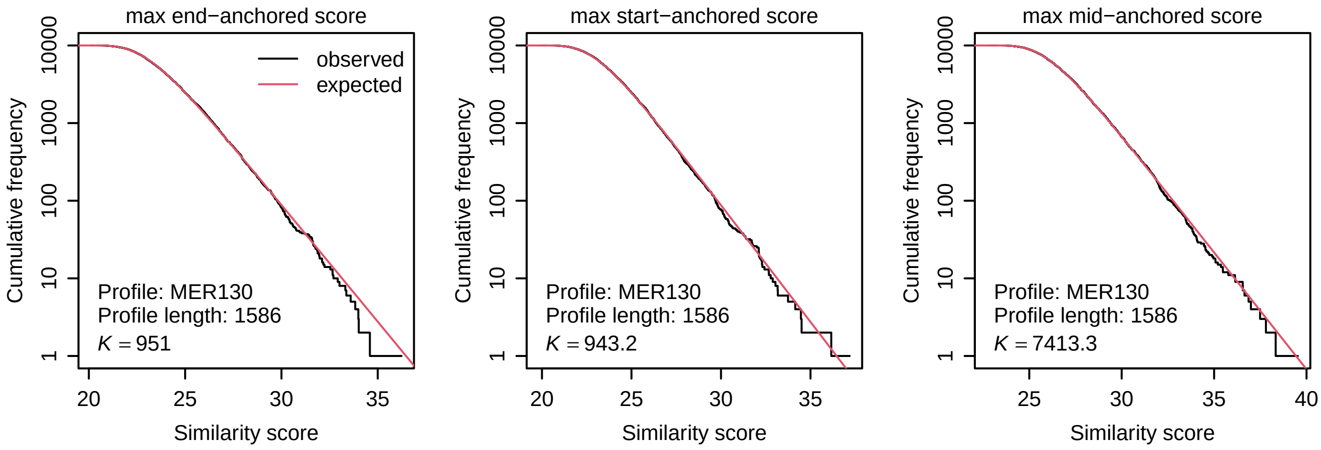


**Figure S11:** Distributions of similarity scores between MER130 (DF000000726.5) and random DNA sequences.

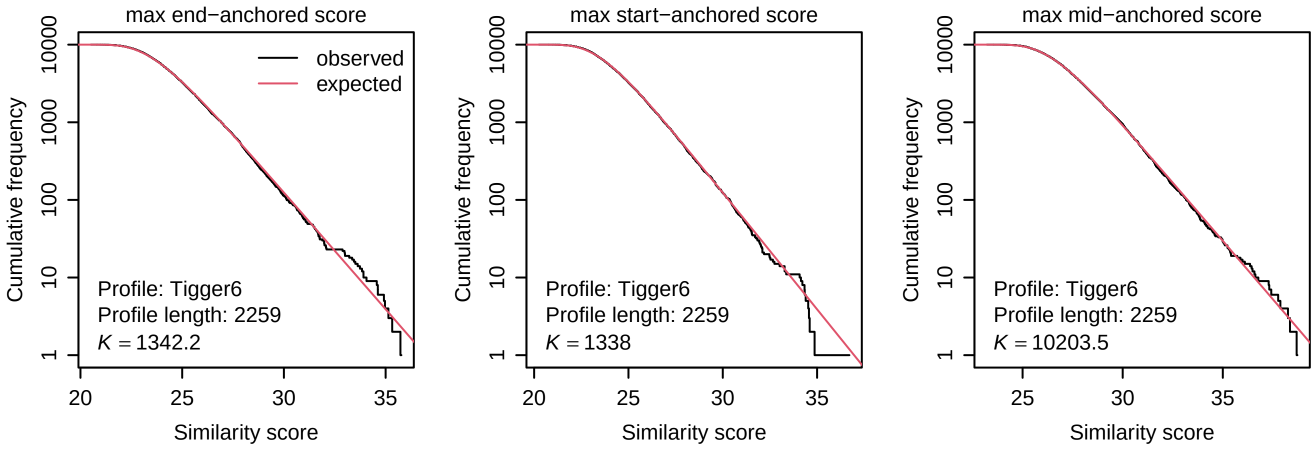


**Figure S12:** Distributions of similarity scores between Tigger6 (DF000001299.2) and random DNA sequences.

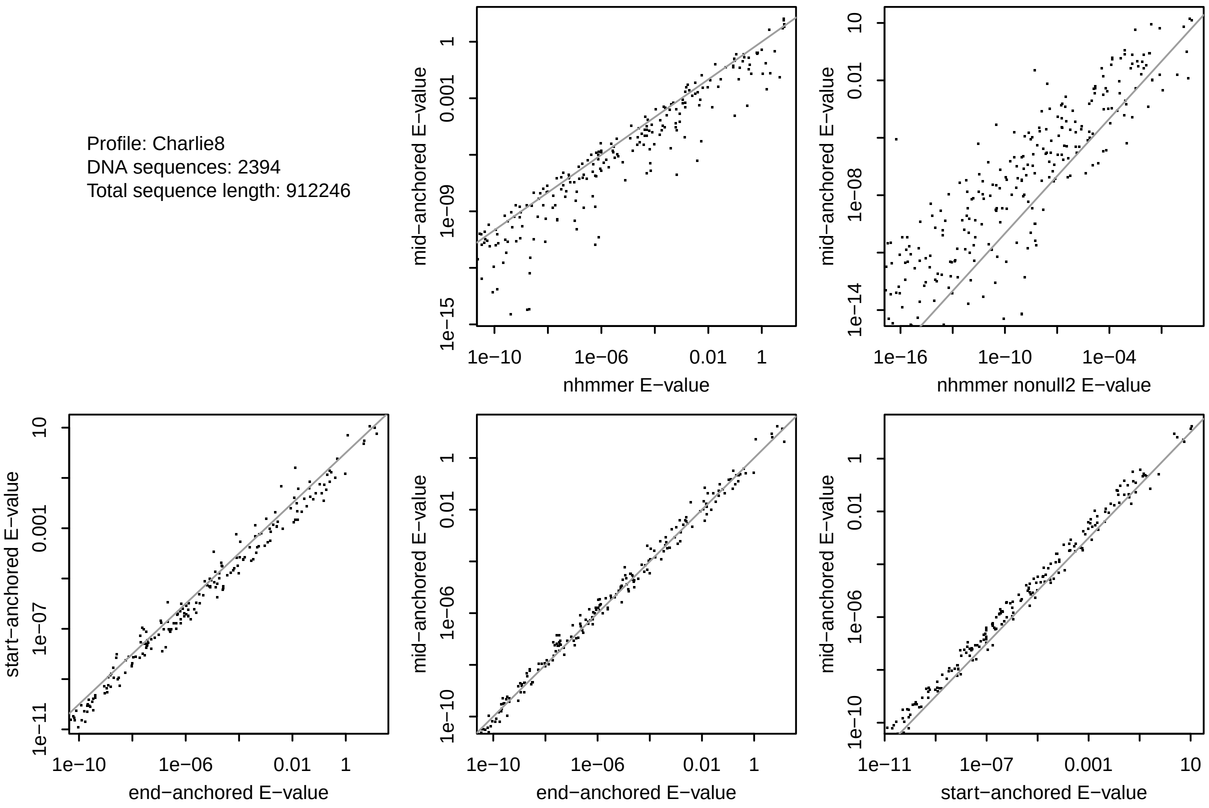


**Figure S13:** $E$-values for similarity between: the Charlie8 profile, and human DNA sequences with Charlie8 RepeatMasker annotation. Each dot shows one DNA sequence. Each sequence has 5 kinds of $E$-value: end-anchored, start-anchored, mid-anchored, nhmmer, and nhmmer with option `--nonull2`.

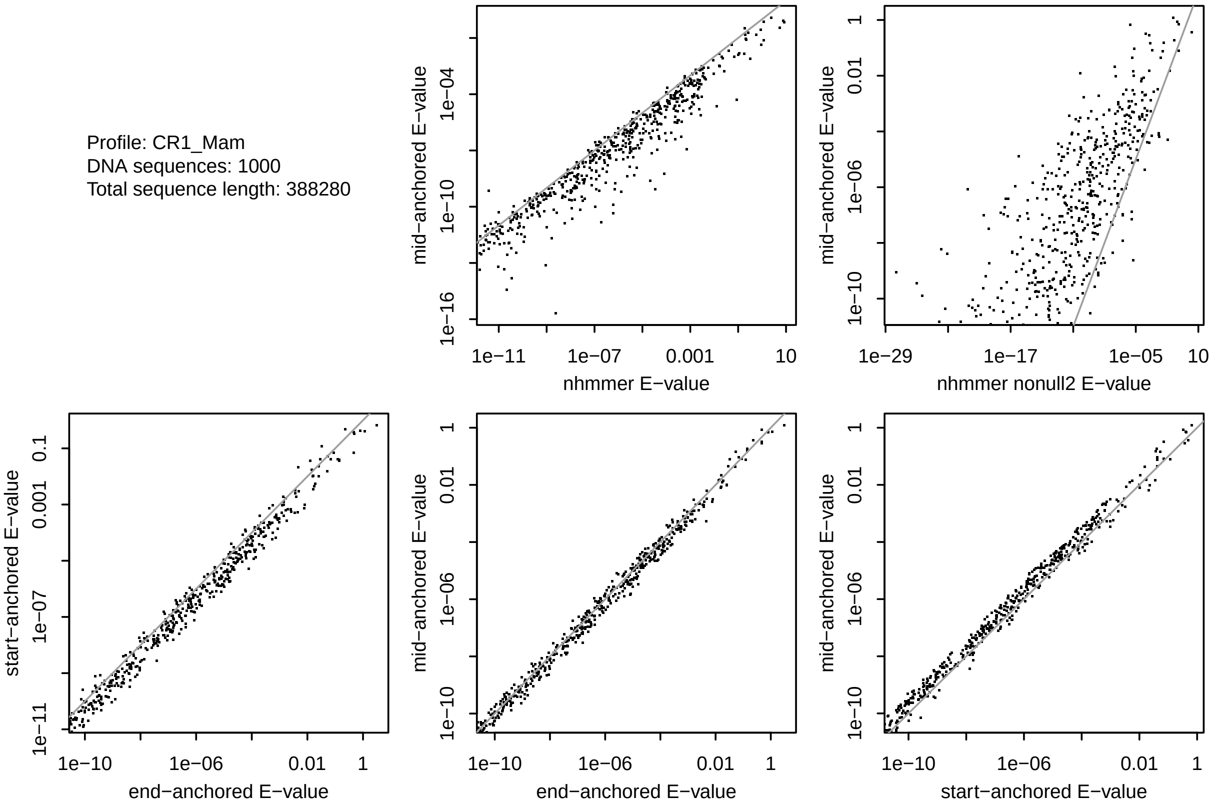


**Figure S14:** $E$-values for similarity between: the CR1_Mam profile, and human DNA with CR1_Mam RepeatMasker annotation.

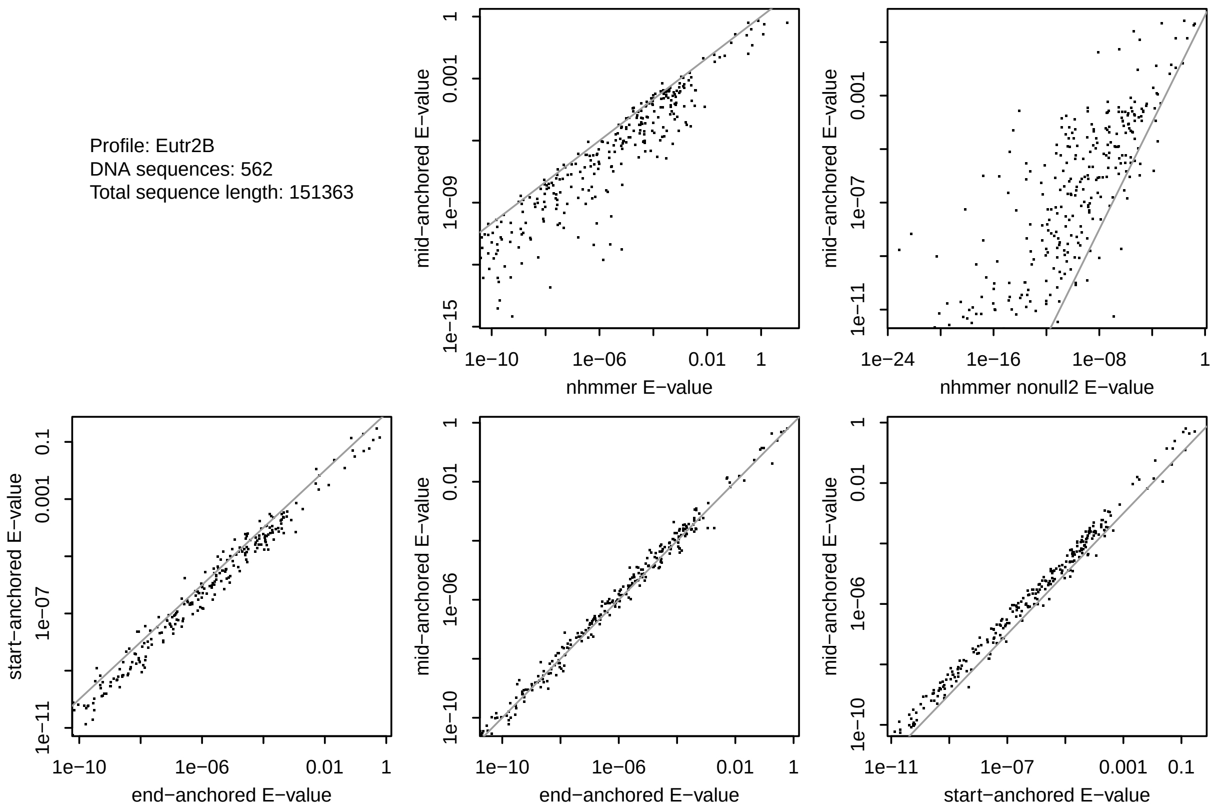


**Figure S15:** $E$-values for similarity between: the Eutr2B profile, and human DNA with Eutr2B RepeatMasker annotation.

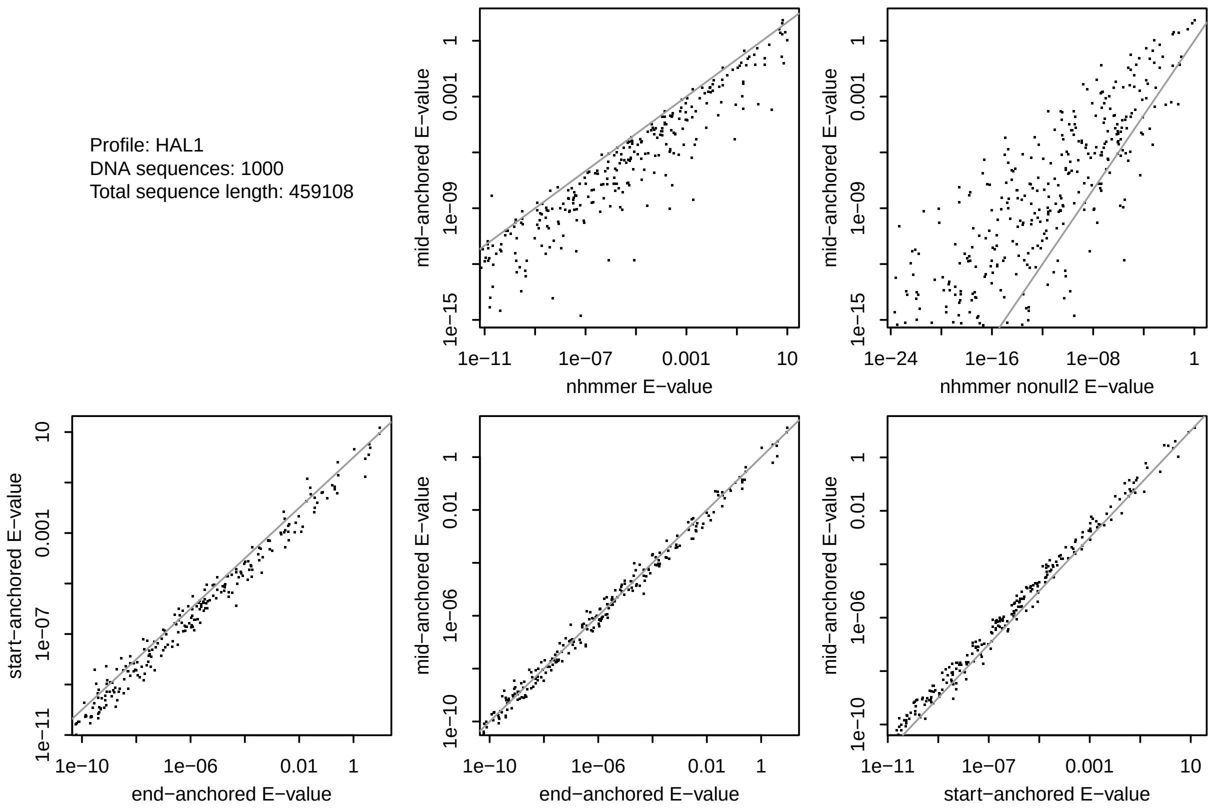


**Figure S16:** $E$-values for similarity between: the HAL1 profile, and human DNA with HAL1 RepeatMasker annotation.

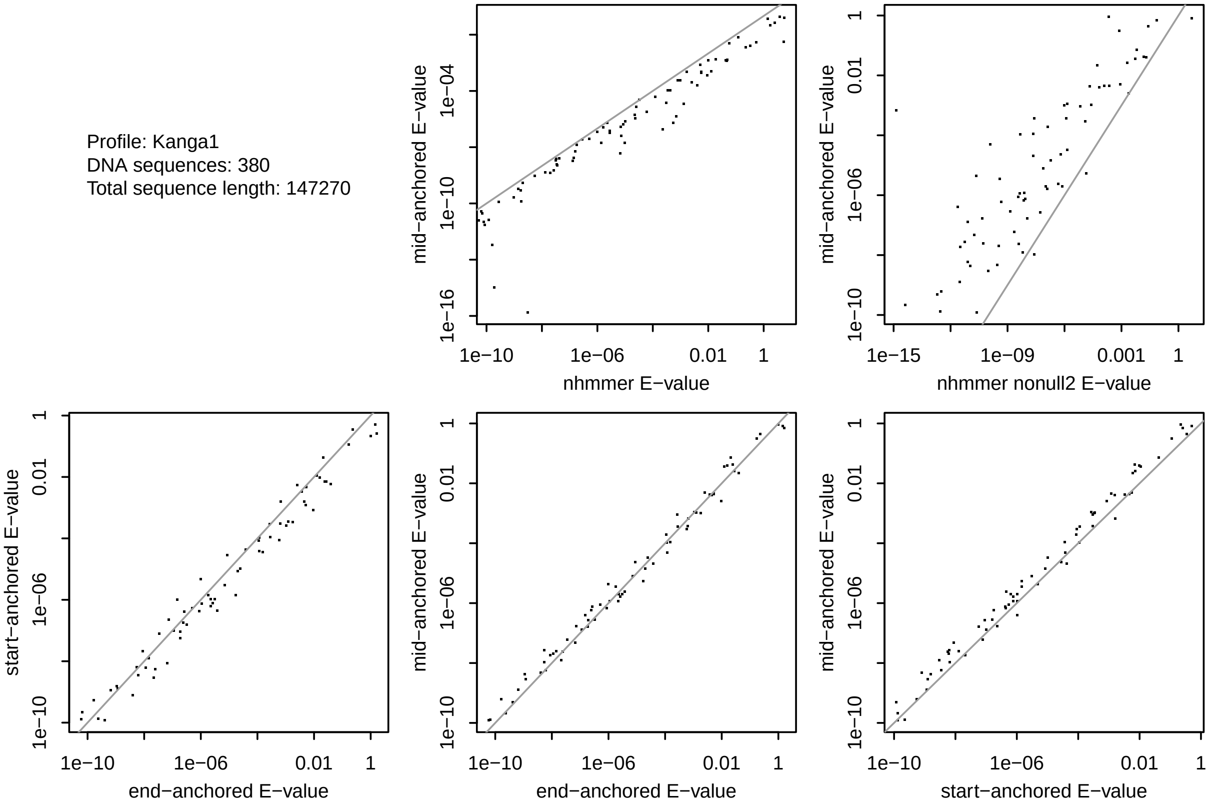


**Figure S17:** $E$-values for similarity between: the Kanga1 profile, and human DNA with Kanga1 RepeatMasker annotation.

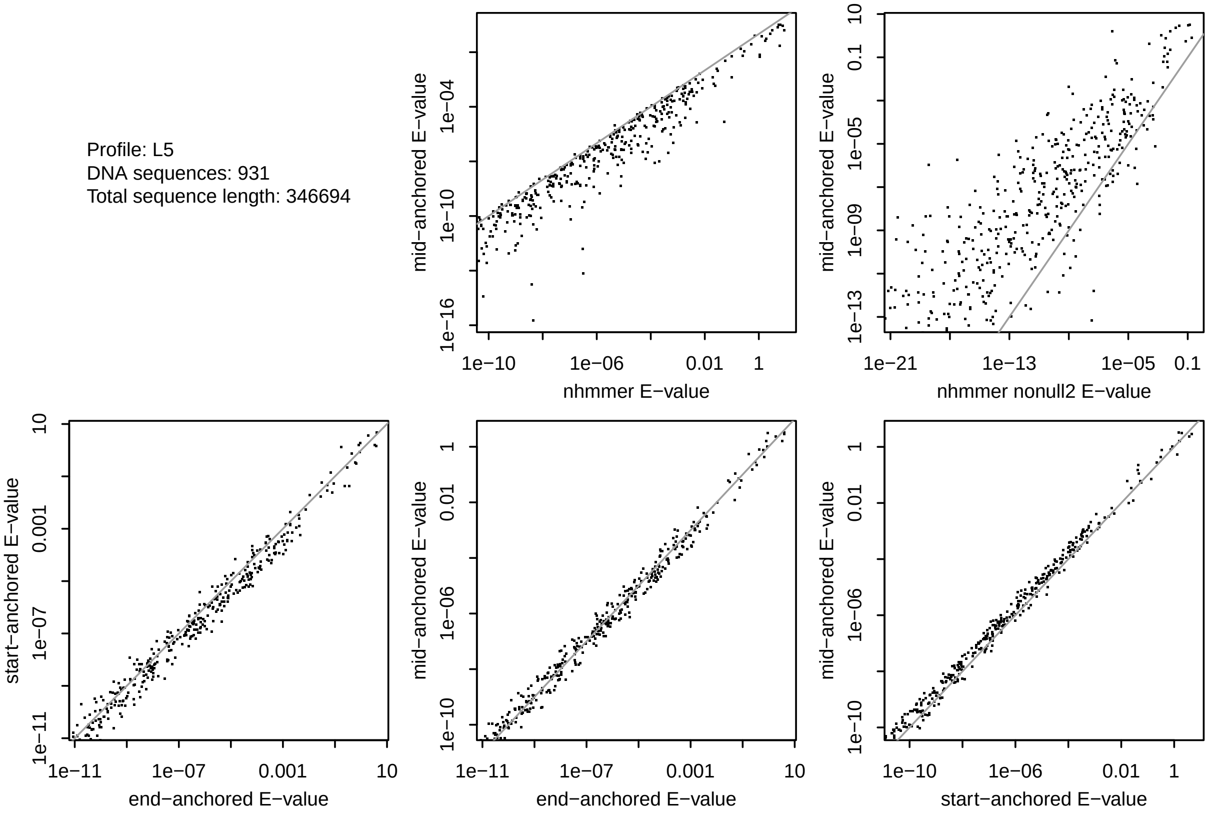


**Figure S18:** $E$-values for similarity between: the L5 profile, and human DNA with L5 RepeatMasker annotation.

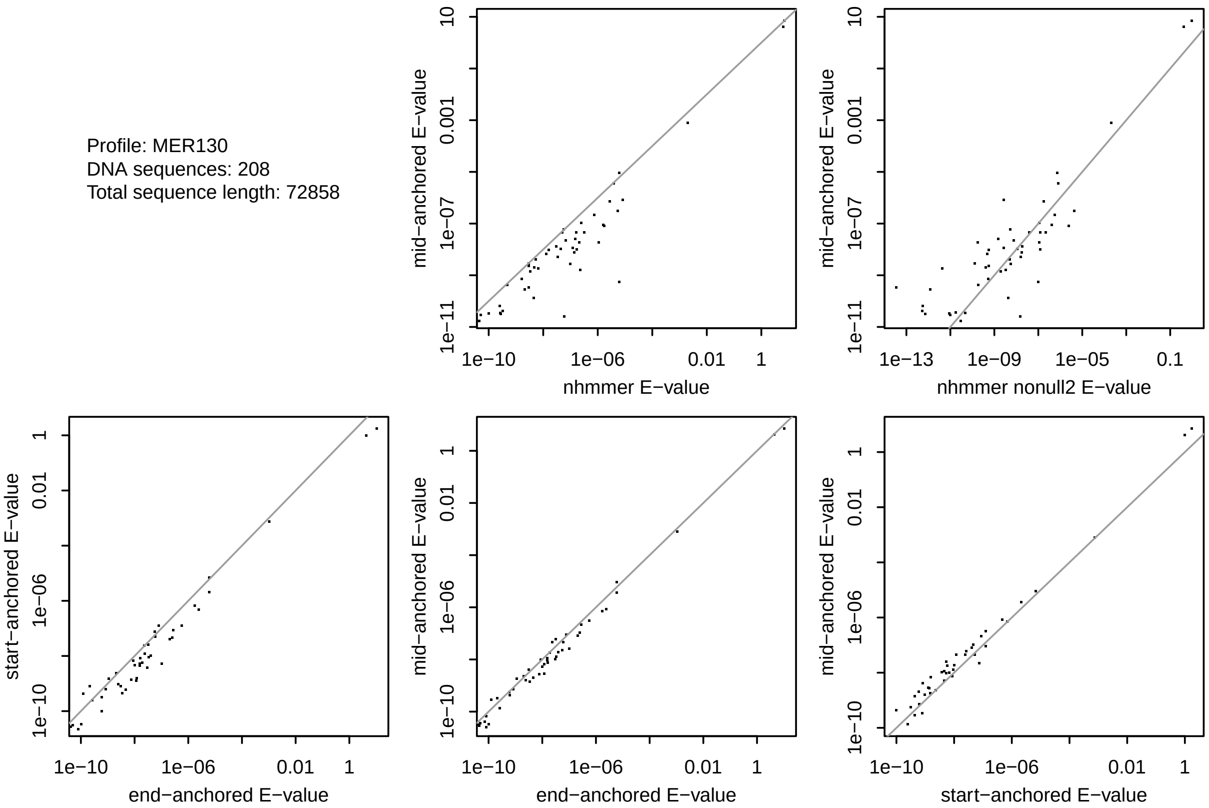


**Figure S19:** *E*-values for similarity between: the MER130 profile, and human DNA with MER130 RepeatMasker annotation.

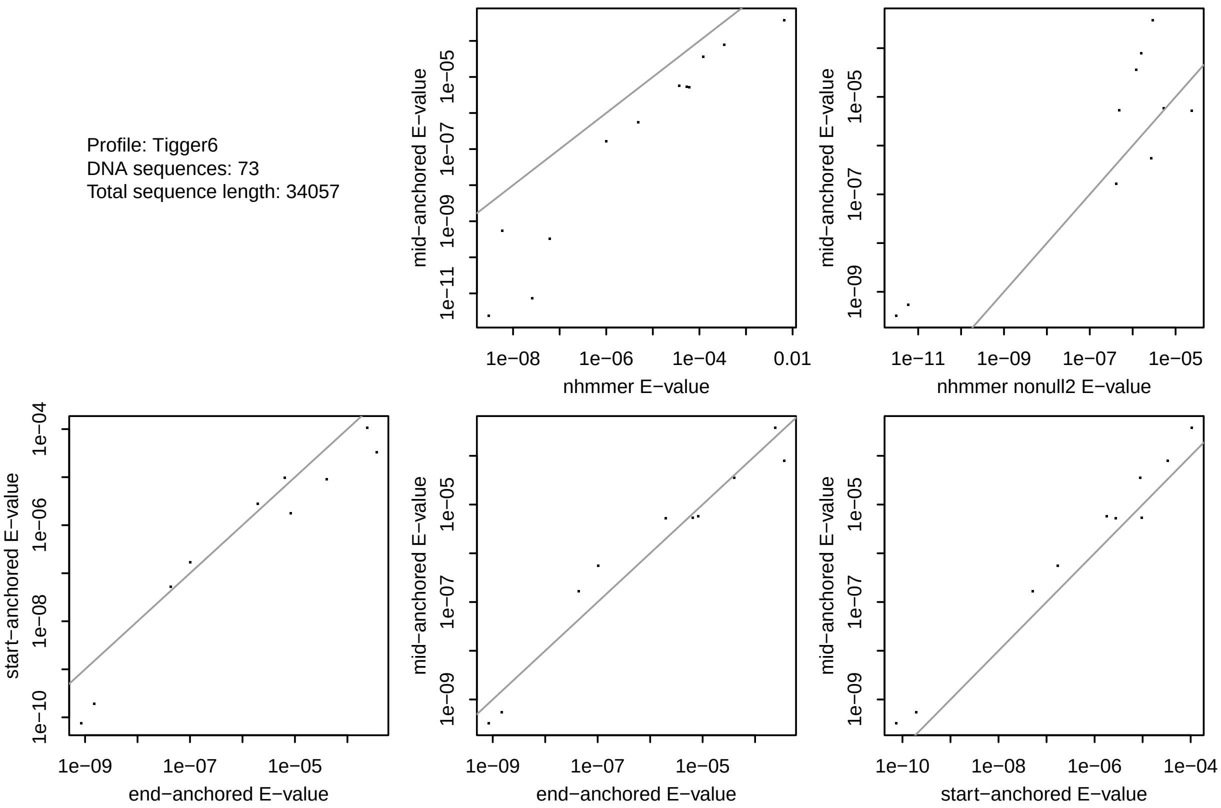


**Figure S20:** *E*-values for similarity between: the Tigger6 profile, and human DNA with Tigger6 RepeatMasker annotation.
